# Images in red: A methodological and integrative approach for the usage of Near-infrared Hyperspectral Imaging (NIR-HSI) on collection specimens of Orthoptera (Insecta)

**DOI:** 10.1101/2024.10.12.617997

**Authors:** Gustavo Costa Tavares, Neirivaldo Cavalcante da Silva, Ana Lúcia Nunes Gutjahr, José Antônio Marin Fernandes

## Abstract

Anthropogenic actions have caused severe environmental changes, leading to a rapid and massive loss of biodiversity, called the Holocene Extinction or "Sixth Extinction." In this context, integrative methods, capable of quickly and reliably determining and differentiating species, have become constantly demanded in taxonomic studies. Near-infrared spectroscopical technologies have proved to be promising for application in the integrative taxonomy of insects, and it is a true-non-destructive method, causing no damage to the sample and being completely chemical-preparation-free. Near-infrared spectral profiles are known as the fingerprints of the chemical composition of a given sample, and a new layer of information may be accessed. Hyperspectral imaging technologies in the near-infrared range are among the most popular, although their usage is still incipient in insect studies. As in other animal taxa, katydids’ taxonomy has traditionally been based on morphological comparison, resulting in many misclassifications over the years. However, integrative methods are more and more required in taxonomic studies. Different methods and technologies have been used from an integrative perspective to minimize misidentifications, especially for non-taxonomist or untrained researchers. Here, we approach the applicability of near-infrared spectroscopy coupled with hyperspectral imaging technology to Ensifera specimens housed in collections, discussing the advantages, disadvantages, and challenges to future applications. As a result, we propose using only homologous body parts in comparing and modeling species using this kind of data due to the heterogeneity of the insects’ exoskeleton. Additionally, we made a case study by discriminating four species of the katydid genus Conocephalus Thunberg, one of the most speciose genera within Orthoptera that is known to have polymorphic species, with variations expressed in the wing development and postabdomen appendages. We generated a Partial Least Squares-Discriminant Analysis (PLS-DA) classification model for the species with an overall classification accuracy for the assigned pixels of 90% and specific accuracy for pixel discrimination ranging from 96% to 98%. This is one of the few research studies employing hyperspectral imaging in insects’ taxonomy, and it is the very first to use this technology in Orthoptera; therefore, this is a preliminary approach to usage as an integrative method.

## Introduction

Anthropogenic actions have caused severe environmental changes, leading to a rapid and massive loss of biodiversity, which has been called the Holocene Extinction or "Sixth Extinction" [1,2]. It is estimated that more than half of the terrestrial surface of our planet is altered, causing an average extinction rate of 25000 species per year [3,4]. This catastrophic scenario adds a sense of urgency to the work of taxonomists, who have provided universal names and classification systems of biodiversity for almost three hundred years [5,6]. This sense of urgency is more evident when we face three other alarming estimates: (1) approximately 90% of all Eukaryotes are expected to be unknown [3]; (2) the average time between the collection of a specimen of a new species and its formal description (time that it is stored in a collection waiting for a specialist) is 21 years; and (3) if this rate of description is maintained, it will take at least another 10000 years to describe all the current fauna and flora [3,4,7].

In the last two decades, many scientists have suggested that the traditional taxonomy is in crisis, called taxonomic impediment. This crisis is driven by the dramatic environmental changes caused by human activities, the rapid loss of biodiversity, the small and decreasing number of taxonomists, the long delay in species descriptions, the instability of name validity, and the credibility of descriptions and classifications based only on morphological comparison [3,6,16– 20,8–15]. In a proposal for an approach to taxonomic practices, Dayrat [21] coined the term integrative taxonomy. For him, integrative taxonomy aims to delimit the units of the diversity of life from multiple and complementary perspectives. In other words, a species must be delimited under a multidisciplinary approach, with information from not only morphological comparison but also, for instance, phylogeny, phylogeography, population genetics, ecology, ethology, etc. In this sense, the search for new methodological and technological tools that allow a fast and reliable characterization and that add more information complementary to the morphology of the different forms of life on Earth is constant, especially when facing a taxonomic group as diverse as insects [6,8,22–24].

Molecular methods have mainly been used as a reliable methodology for complementary data in integrative taxonomy, with DNA barcoding being the most notorious and widely used molecular method [25–27]. The mitochondrial gene Cytochrome Oxidase I (COI) has been proposed as the center of a global identification system for animals, the BOLD project (The Barcode of Life Data System) [28]. However, museum specimens’ DNA sequences are usually highly degraded due to many aspects involved in the post-mortem preservation and age of the stored specimens, such as using chemicals, humidity, temperature, and softening processes [29,30]. Although some recent DNA extraction methods are called "nondestructive" [29,30], they employ the usage of lysis buffer and proteinase K, which preserves the mechanical structure of the exoskeleton but digests all the internal components and alters the cuticle composition, making it completely hyaline.

Metabolomics can provide helpful information for integrative taxonomy using collection specimens since it is the study of the metabolites or global characterization of metabolic molecules in an organism [23,31–34]. Metabolomics is one of the areas of science used in chemotaxonomy, the biological organization of organisms based on their chemical markers, which is already widely exploited in plant taxonomy [35–37]. However, conventional metabolomic methods are generally expensive, time-consuming, or destructive since they may involve chemical preparation, sample mechanical powdering, and exposition to elevated temperatures or more energetic radiations of the electromagnetic spectrum. On the other hand, Near-infrared spectroscopy (NIRS) is a lower-cost alternative that almost instantly provides massive information about a sample’s chemical composition [38–41].

NIRS is an emerging nondestructive analytical technique based on the region of the electromagnetic spectrum between 750 and 2500 nm, which can change the stage of molecular vibrational energy and thus obtain, as a spectrum, qualitative and quantitative information from the unique interaction between different molecules and near-infrared radiation [23,34,42]. Besides the technique not destroying the sample, it also has other advantages, such as the fast acquisition of spectra, non-generation of residue, and no need for sample chemical preparation [34,42,43].

The applicability of NIRS is vast and promising in many fields of entomology [40]. Regarding the usage in insect taxonomy, the technique can distinguish taxa from intra- and interspecific levels to higher taxonomic ranks, such as genera, families, or orders [39], and has been tested in pinned [44] and alcohol-preserved museum specimens [45]. A very recent example was reported by Johnson [46]. He successfully distinguished three genera of weevil beetles with an accuracy rate ranging from 92 to 96%. The method’s efficiency has also been explored in forensic entomology by differentiating maggot and adult fly species that visit carcasses [47–49].

Although the literature shows how the method is promising, it is essential to bear in mind that, in order to be widely applicable in insect taxonomy, the spectrum acquisition method must be standardized so that the data obtained may be comparable between the different taxa. In this sense, the method applied to collect data for a given taxonomic level must be repeatable for the others. However, each author adapted the methodology for spectrum acquisition, using entire specimens [41,45], powdering the specimen [50], or only detached body parts [44,51].

Another point to be considered is that cuticle formation is the last step of an insect’s embryo development, so any alteration in the DNA would affect the cuticular composition, and different body parts of an insect (i.e., non-homologous parts) have a distinct cuticular composition [52]. Considering the high detection sensitivity of NIRS, these differences probably result in different spectra profiles, even if subtle. One way to overcome the size limitations and consider cuticle heterogeneity, especially in separately analyzing non-homologous regions, is to use hyperspectral imaging (HSI) equipment operating in the near-infrared (NIR) range.

HSI is a particular type of image in which a continuous spectrum is acquired for each pixel [53]. These images form a data structure known as a hypercube, where each pixel contains a spectral measurement captured by hundreds of channels, each corresponding to a specific region of the electromagnetic spectrum. This differs from standard digital cameras, which capture images using only three channels—red, green, and blue (RGB) [54]. So, HSI is a visual representation of a sample as a function of a specific region of the electromagnetic radiation spectrum. When these images are taken in the NIR range, this ensures the analysis of the chemical components on a given surface and the visualization of how they are distributed [55]. Among the spectroscopic analytical techniques combined with imaging technologies, NIRS has been very promising and is among the most popular [53,55–57]. It is a high-speed method that can be perfectly usable for integrative taxonomy once it almost instantly provides an enormous amount of information. The nondestructive characteristic may be the best alternative for museum specimens, especially when leading with types. However, it is worth mentioning that the hypercubes are very large files, and handling them requires computers with high processing capabilities once the three-dimensional cubes have an enormous quantity of data [58].

In entomology, HSI is usually used to detect damage to cultivars or contamination by pests, with spectra in the visible range and/or part of the NIR being most often used [56,59–65]. Despite the incipient usage in insect taxonomy, Wang et al. [66] made the first attempt at the usage of NIR-HSI (with spectra ranging from the visible to NIR) from an integrative perspective, also combining morphological and DNA barcoding data, to successfully differentiate seven lineages of the leafhopper genus *Bundera* Distant.

Only two works apply NIRS techniques to Orthoptera species [44,51], but none use HSI technologies. Grace et al. [51] acquired spectra from the pronotal disk of two populations of the grasshopper *Hesperotettix viridis* (Thomas) using bands in the visible spectrum and NIR (450-2500 nm). The authors reported that the two populations (which feed on different plants) significantly differ in cuticular lipid composition. In Sovano et al. [44], the authors revalidated the name *Aganacris sphex* (Rehn) based on NIR spectra, being the first work in which a nomenclatural act was made based on NIR spectra. Katydids of the genus *Aganacris* have a high sexual dimorphism, and males and females were historically described as different species [67]. However, Sovano et al. [44] reported that they were able to group males and females of all species based on NIR spectra.

Morphological characterization has traditionally defined taxonomy in katydids, resulting in some controversial classifications [68,69]. Mugleston et al. [70–72] have suggested that this concept has led to misclassification, and many subfamilies and tribes are, in fact, paraphyletic. As a result, many genera are hard to assign to tribes and are still *insertae sedis* [73].

*Conocephalus* Thunberg is one of the most taxonomically challenging katydids genera and the genotype of the subfamily Conocephalinae. Its representatives are distributed worldwide and are generally small greenish katydids inhabiting grasslands and other open areas [73,74]. It is one of the most diverse genera in Tettigoniidae, with 167 valid species (and 647 invalid specific names) occurring in almost all continents except Antarctica [73]. Despite being quite common, only 11 species are recorded in Brazil, with few official records [73,74].

Historically, species’ differentiation in *Conocephalus* relies mainly on the male external genitalia. However, many species are polymorphic, with variations in wing development or genitalia, resulting in species with two or three described cerci shapes known (e.g., *Conocephalus saltator* (Saussure, 1859) and *Conocephalus versicolor* (Redtenbacher, 1891)) [75–78]. Females may also present polymorphisms in postabdomen appendages, with the ovipositor varying in length and shape [76,78], which may make it difficult to determine the species in the absence of a male specimen.

So, this work considers all the context and issues of the traditional taxonomy, the increasing necessity of new methodological approaches to be applied in integrative taxonomy, the operational and analytical advantages of hyperspectral imaging in the near-infrared range, and all the historical taxonomic problems of the order Orthoptera, especially the genus *Conocephalus*. In this sense, this work aims to approach the methodological applicability (advantages and disadvantages) of near-infrared spectroscopy coupled with hyperspectral imaging technology to Orthoptera specimens housed in collections as an innovative method for integrative taxonomy. It also aims to test the efficiency of this technology in discriminating species, using four species of the katydid genus *Conocephalus* as a model.

## Material and methods

### Samples preparation

We organized our samples into two analytical sets. The first set was used to explore the advantages and limitations of NIR-HSI in dry specimens; the second set was used to test the efficiency of this technique in discriminating specimens, using four species of the katydid genus *Conocephalus* as a model.

In the first set, we used 143 Orthoptera specimens belonging to three families of the suborder Ensifera and eight subfamilies: Conocephalinae (n=110), Listroscelidinae (n=2), Pseudophyllinae (n=14), Meconematinae (n=7), Phaneropterinae (n=7), and Pterochrozinae (n=1) belonging to Tettigoniidae, Gryllacridinae (n=1) belonging to Gryllacrididae, and Lutosinae (n=1) belonging to Anostostomatidae (S1 Table). All the specimens are pinned and dried and are housed in the Coleção Zoológica Didático-Científica Dr. Joaquim Adis of the Universidade do Estado do Pará (CZDC-UEPA), laboratory of Invertebrates of Universidade Federal do Pará (LA-INV), Museu Paraense Emílio Goeldi (MPEG), and Coleção Zoológica da Universidade Federal do Mato Grosso do Sul (ZUFMS). Each specimen received a voucher label with a unique code (S1 Table). The specimens were gathered from various expeditions, subjected to different killing and preservation methods, and exhibited varying states of conservation.

In the second set, we used only 48 specimens of the original set, belonging to four species of the katydid genus *Conocephalus* Thunberg: *Conocephalus fasciatus* (De Geer, 1773) [males=10, females=4], *Conocephalus versicolor* (Redtenbacher, 1891) [males=6, females=3], *Conocephalus iriodes* Rehn & Hebard, 1915 [males=8, females=2], *Conocephalus saltator* (Saussure, 1859) [males=8, females=9], and *Conocephalus equatorialis* (Giglio-Tos, 1898) [males=2, females=2].

The specimens were fixed on rectangular styrofoam plates measuring 15 x 8 x 0.5 cm covered with a sheet of black cardboard and then leveled so that their dorsal surface was approximately at the same height (Fig 1A, 1B, S1 Fig). The samples were arranged in rows to maximize the scanning process (Fig 1C). Longitudinal lines, 1 cm spaced, were made on the cardboard to mark the space where the samples were fixed (Fig 1C). Plates made of any material that allows a pin to be fixed can be used. We chose styrofoam because it is an easily found and low-cost material, and the pinned specimens can be fixed to be scanned.

**Fig 1.**
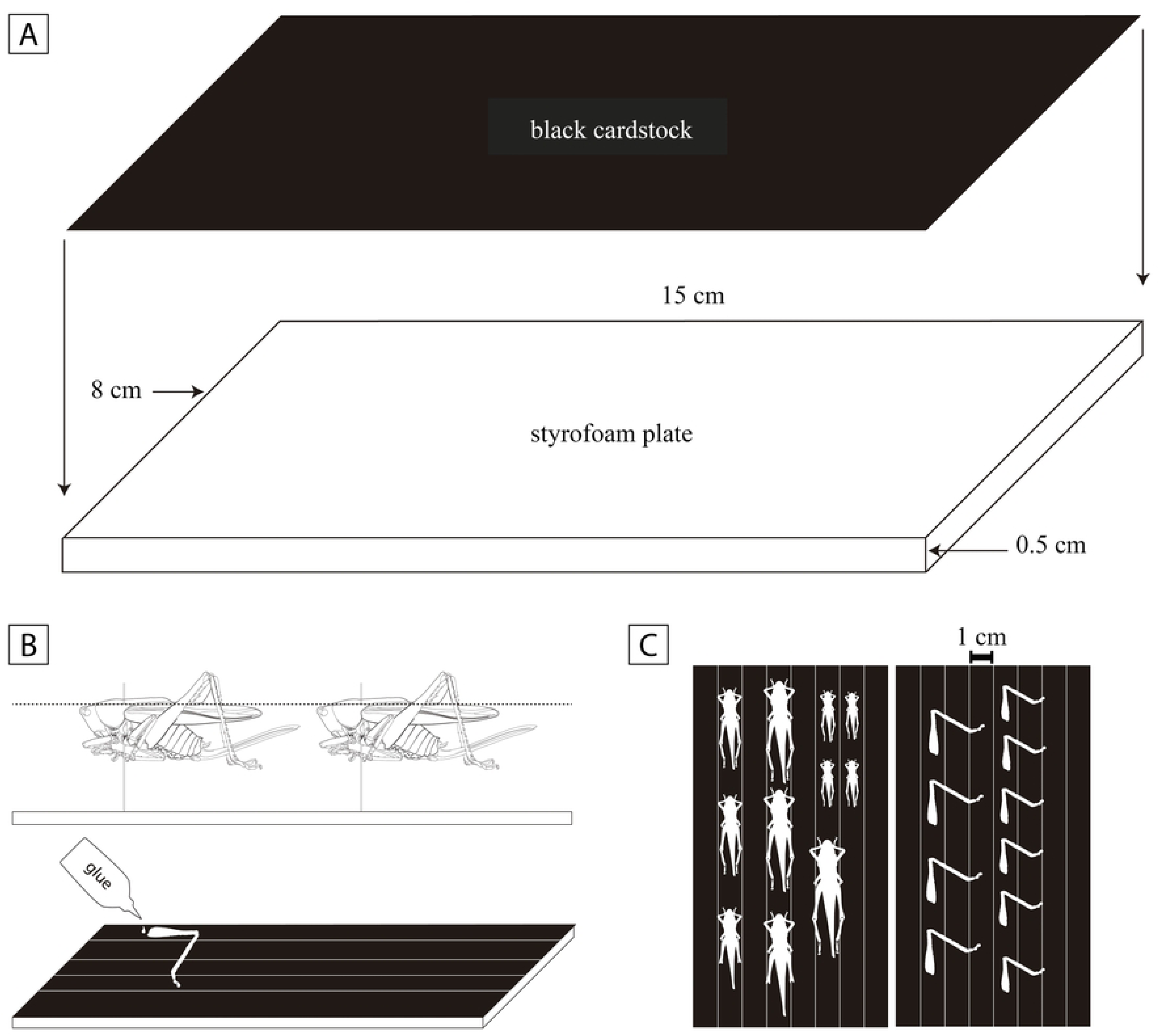
Image acquisition protocol for dried museum specimens. (A) Styrofoam plate confection. (B) Specimens leveled to the same height and legs glued to the styrofoam plates. (C) Samples organization for scanning.

One hind leg was removed from each specimen. A tiny drop of glue was used on the base of the detached hind legs to affix them to the confectioned plates, with their external surface exposed (Fig 1B) and organized in rows (Fig 1C). Gluing the legs is not mandatory. The hind legs may be arranged directly on the equipment’s Teflon plate. However, we chose to glue the legs on the same confectioned plates to [1] compare both images of the body and legs under the same background influence and [2] to evaluate the penetration capacity (transmittance) of near-infrared radiation.

### Hyperspectral imaging acquisition

The images were acquired with a Hyperspectral Imaging Spectrometer Sisuchema-SWIR (manufacturer: Specim, Spectral Imaging Ltd., Oulu, Finland). We used the software ChemaDaq (version 3.621.992.6-R) for image acquisitions. A 10 mm macro lens was used, with a spatial resolution of 31.25 µm for each pixel with a maximum image width of 1 cm (320 pixels). These images were taken in reflectance measurement mode and scanning spectral acquisition mode, with a spectral range between 900 and 2493.75 nm (256 channels) and a spectral resolution of 6.25 nm. The frequency was adjusted to 60 Hz and the exposition to 2 ms.

The focus was adjusted to emphasize the head and pronotal disc for the full-body images and the femur’s outer surface for the images of the hind legs. Three images of each sample were made (triplicate) in slightly different focus positions to minimize the chances of error and test the focus’s influence on the results. The temperature during the image acquisition varied from 19 to 20.1 °C, and the air humidity from 51 to 73% (S2 Table).

### Pre-processing

All the images and spectral treatment were made with MATLAB (MathWorks, Inc., Natick, MA). Evince software (Prediktera AB) was utilized to convert the images from .raw format to .mat format (MATLAB compatible format). No spikes or dead pixels were removed in Evince once we could not set the parameters of the removal algorithm. Exploratory analyses and modeling were conducted utilizing HYPER-Tools (version 3.0) [53], the MATLAB Free Graphical User Interface (GUI) for Multispectral and Hyperspectral Image Analysis.

Spectra with at least 25% of zeros or missing values were considered dead pixels, following Vidal and Amigo [79]. Moboaraki and Amigo [53] recommended that a factor of six times the standard deviation of the average spectrum should be used as a threshold to detect and eliminate spikes of a hyperspectral image. However, different thresholds were tested in the first set of images, progressively reducing the value from six to one (i.e., equal to the standard deviation mean spectrum) to verify the capacity to remove spikes in our samples. The threshold values that better worked in the first set were applied to the second set.

For the first set, cropping and spatial binning of the images were performed using two and three window sizes to reduce calculation times and minimize ratio noise. The pronotal disc and the hind femur’s proximal half were chosen as regions of interest because they were the areas in focus with fewer irregularities on the surface, aiming to avoid radiation scattering. For spectral preprocessing, Savitzky-Golay polynomial derivative and smoothing filter [80] with a first-order polynomial (2^nd^ degree) with window size varying from 9 to 21 and Standard Normal Variate (SNV) [81] were tested to enhance signal-to-noise ratio and minimize radiation scattering. Potential edge-effects were reduced by excluding variables at the ends of preprocessed spectra, according to the window size of the Savitzky-Golay smoothing filter applied. The k-means method, with two or three clusters, was employed to remove the background when necessary. Additional manual removal was made when required.

For the second set, only the pronotal disc was used, employing a window size of 2 (2x2 pixels²) for spatial binning. Spectral preprocessing and exclusion of edging wavelengths were performed using the approach that worked better for the first set.

### Chemometrics and spectral profiling

For the first set, prospective analyses were made using Principal Components Analysis (PCA), with the purpose of finding congruent and divergent spectral patterns between samples. PCA is an exploratory method used to visualize and analyze complex datasets by finding the directions in the data that explain the maximum variance. Such directions are termed Principal Components. The coordinates of the original data points (pixels in this case) in the new coordinate system defined by the principal components are called scores, and the magnitude of influence of each original variable (NIR wavelengths in this case) on the principal component are called loadings. While the scores provide insights into how each sample contributes to the overall variation in the dataset, the loadings are crucial for interpreting the underlying structure of the data and identifying the variables that drive the observed patterns [82]. The PCA was performed in paired images, comparing the spectrum of the legs and pronotum of the same specimen and homologous regions of different individuals. When necessary, specific areas of the hypercubes were unfolded to enable a more detailed analysis of the spectra.

For the second set, triplicate images of five specimens from each species of *Conocephalus* were utilized (S2 Table and S2 Fig), except for *C. equatorialis*, for which only four specimens were available. A preliminary PCA prospective analysis was also performed in this case. A Partial Least Squares Discriminant Analysis (PLS-DA) model was employed to classify species of *Conocephalus.* PLS-DA [83] works by establishing a latent variable (LV) structure aimed at capturing the maximum covariance between the predictor variables (NIR wavelengths in this study) and the categorical response variable, representing the species classes of *Conocephalus* (*C. fasciatus*, *C. versicolor*, *C. iriodes*, *C. saltator*). Accordingly, PLS-DA effectively seeks to segregate pixels belonging to each class within the latent variable space. During the training phase, individual modeling of each class is conducted, assigning a binary value of 1 to the respective class and 0 to all other classes. Given that the PLS-DA classification method utilizes partial least squares regression (PLS), it results in continuous categorical variable modeling. Consequently, predicted values are not strictly bounded to the binary scale of 0 or 1, making it necessary to establish a threshold value as a discriminant criterion. The Bayes decision rule was employed to determine this threshold. *C. equatorialis* was not included due to the small number of its specimens. Venetian Blinds Cross-validation (with samples divided into ten groups) was utilized for the PLS-DA models.

The optimal number of latent variables was chosen based on the analytical performance of the PLS-DA classification evaluated using sensitivity (Sn) and specificity (Sp) metrics. Sensitivity indicates the model’s ability to accurately classify samples belonging to the class of interest, while specificity reflects its ability to correctly reject samples not belonging to the class of interest, as defined by the equations below:

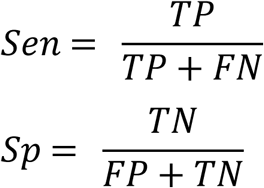

TP, TN, FP, and FN represent the numbers of true positives, true negatives, false positives, and false negatives, respectively. The model was then tested on images of the remaining specimens of *Conocephalus* (including those of *C. equatorialis*) to verify the efficiency in identifying not-modeled specimens. We considered that the model correctly classified an image when 60% or more of the pixels were correctly assigned to the taxa previously identified based on morphology, partially correct when between 30% and 59%, and failed when less than 30%.

Variable selection was not conducted before the PLS-DA modeling stage. Typically, variable selection techniques are employed to improve model performance by identifying a subset of variables that better differentiate sample classes or correlate more strongly with analyte content or relevant properties, functioning as a more targeted form of spectral preprocessing [84]. Nevertheless, we opted against selecting variables in order to evaluate the method’s efficiency under conditions of maximal external interference. Achieving positive results in such a complex matrix would serve as strong evidence of the HIS-NIR method’s effectiveness.

When necessary, False RGB images were created by dividing the spectra of each pixel into three portions with equal numbers of points, calculating the mean value of each portion to generate a representative value for each RGB channel, and normalizing each mean value.

## Results and discussion

### Sample preparation, preservation, and time-consumed

The rectangular styrofoam plates with black cardboard sheets made the scanning samples process much faster. Each round of scanning lasted between 30 and 90 seconds. By using this method, it was possible to scan multiple samples at the same time. Although larger specimens, such as those from the genus *Ceraia* Brunner von Wattenwyl, were scanned individually or in pairs, up to five smaller specimens, like those of *Phlugis* Stål, could be scanned simultaneously (S1 Fig). However, standardizing the focus position across multiple samples can be challenging. Performing triplicate scans increases the likelihood of having all specimens in focus in at least one image, though researchers may opt to scan each specimen individually for better precision.

As Zheng et al. [58] mentioned, HSI images are very large data files, and dealing with them results in a high computational burden. For example, the hind leg of a male *C. fasciatus* is less than 2 cm long, and the entire specimen is slightly longer. However, the individualized image of the hind leg has 160000 pixels, and for the entire body, 192320 pixels, each with a spectral reading captured by 256 channels, meaning 256 columns with the absorbance value of each channel for each pixel (Figs 2A-D). These images are 100 Mb files.

**Fig 2.**
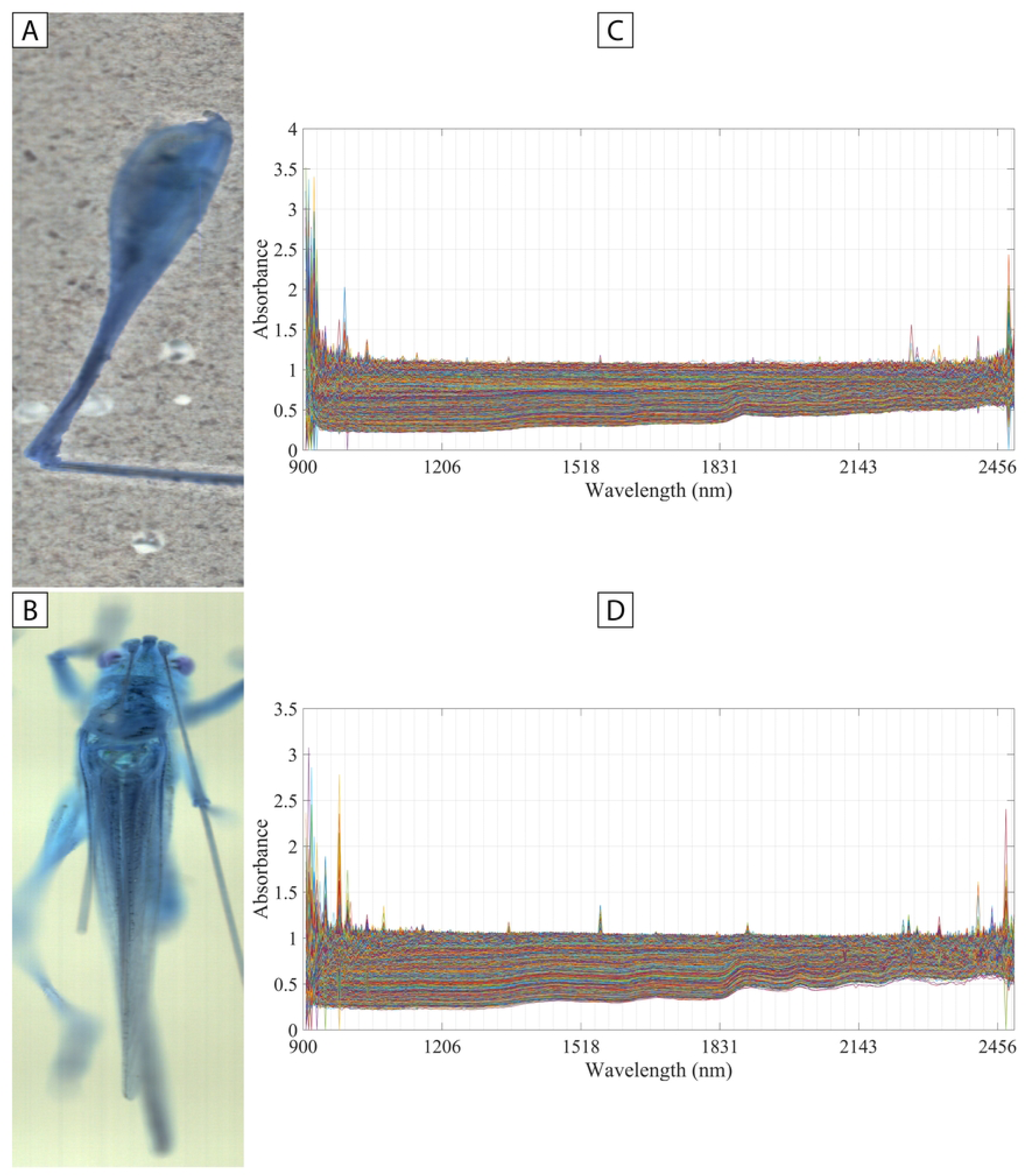
Hyperspectral imaging of a *Conocephalus fasciatus* specimen. False RGB images of (A) a hind leg and (B) body dorsum; (C) 160000 unprocessed spectra of the pixels of figure A. (D) 192320 unprocessed spectra of the pixels of figure B.

### Hyperspectral images

As shown in Fig. 2, the raw full-body images of all specimens contain background spectra and numerous spikes. The background, however, is out of focus and exhibits flat and very noisy spectra with no distinct absorbance bands, making it easily distinguishable from the spectra of the specimens and facilitating its removal during pre-processing. Conversely, the legs, positioned closer to the cardstock, often allow NIR radiation to pass through, resulting in flattened spectra with less pronounced absorbance bands resembling the background (Figs. 2A and 3A-E). The pronotum and leg surfaces, characterized by rough, non-uniform textures, heterogeneous composition, and continuous variations in relief and width, likely cause NIR radiation scattering. This scattering is reflected in baseline fluctuations, which reduce the signal-to-noise ratio [85].

**Fig 3.**
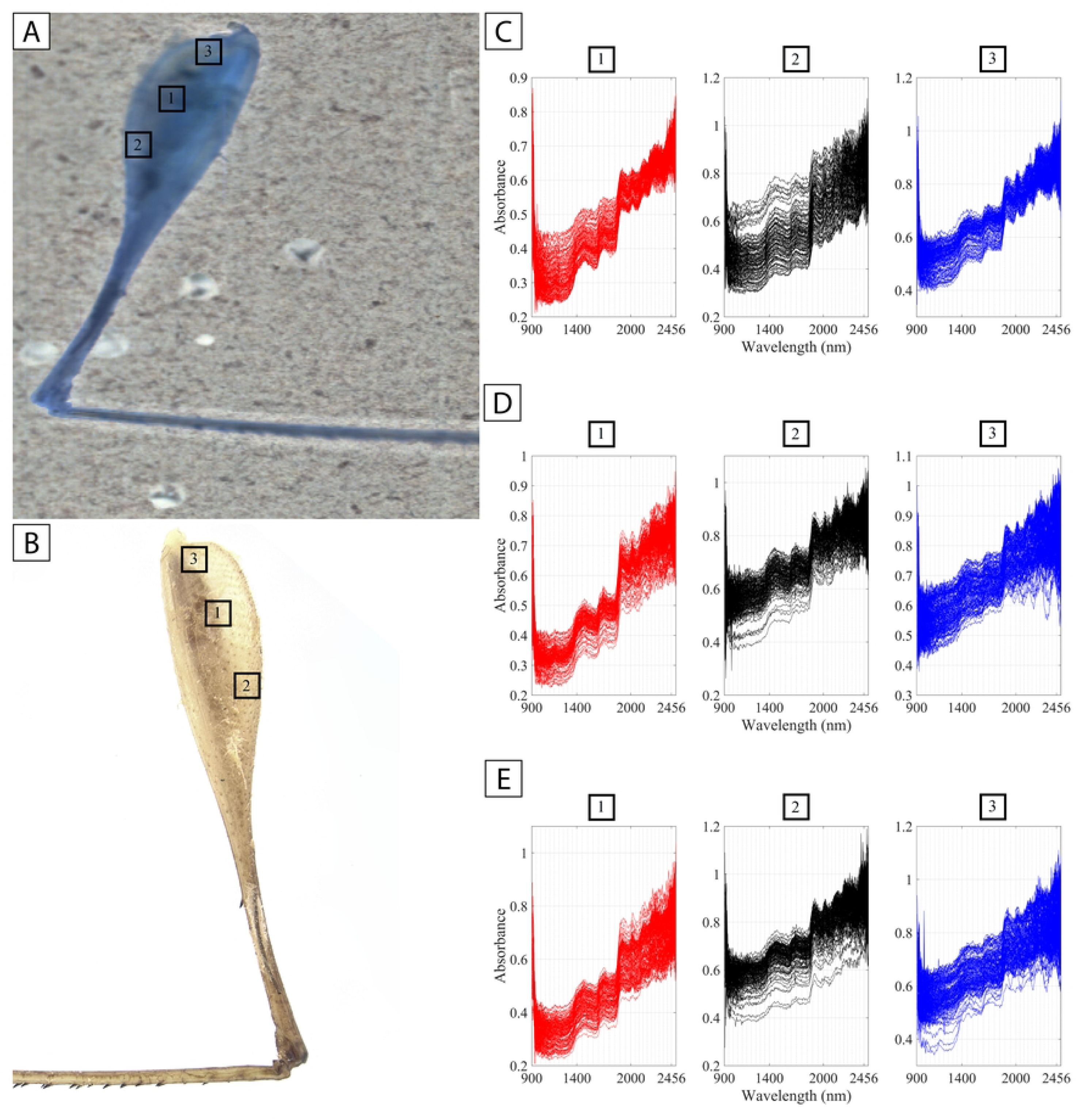
Comparison between the spectra in three different areas and focus depths of a *Conocephalus fasciatus* leg. (A) False RGB image; (B) true RGB image; (C-E) unfold spectra of the three square-marked leg regions in slightly different focus depths.

One notable characteristic of our images is that the regions from 900 to 1093 nm and 2062 to 2493.75 nm tend to exhibit higher noise levels. This noise is difficult to attribute to specific physical or instrumental factors, as it appears to be random. Random noise is generally associated with instrumental and environmental variables, such as electric current fluctuations affecting both the light source and the equipment detector and temperature variations during image acquisition (see S2 Table). Additionally, variations in the composition across the specimen’s surface and in the physical properties of different parts of the animal (such as cuticle, muscle, and skeleton) can also contribute to noise [85]. This latter factor seems to be the most likely cause of the noise observed in our images.

The samples’ spectral profiles are characterized by an absorbance band of low intensity between 1300 and 1600 nm and a more conspicuous one between 1860 and 1950 nm (Figs 3C-E, 4A-H, and 5E). These bands are congruent with many papers regarding the spectral profiling of different insect orders [47,48,51,59,60,86,87] and are probably related to the amide II bond of chitosan and the first overtone band of the amide group stretching, respectively [40,46,88,89]. Undoubtedly, these bands are all arthropods’ signatures (or spectral fingerprints) once they are correlated to chitosan, a natural polysaccharide derived from the deacetylation of chitin [90,91]. According to Johnson [46], this second band could also be attributed to the first overtone of water (Table 1).

**Table 1.** Main absorbance bands present in the samples.

| Spectral range | Functional group |
| --- | --- |
| 1300-1600 nm | Amide II bond of chitosan |
| 1860-1950 nm | First overtone band of the amide group stretching and first overtone of water |
| 1610-1830 nm | CH, CH<sub>2</sub>, or CH<sub>3</sub> first overtones |
| 1970-2060 nm | Amide combination band of chitosan |
| 2060-2180 nm | C-C combination band |
| 2180-2360 nm | CH<sub>2</sub> or CH<sub>3</sub> combination band |
| 2360-2500 nm | CH combination band |

**Fig 4.**
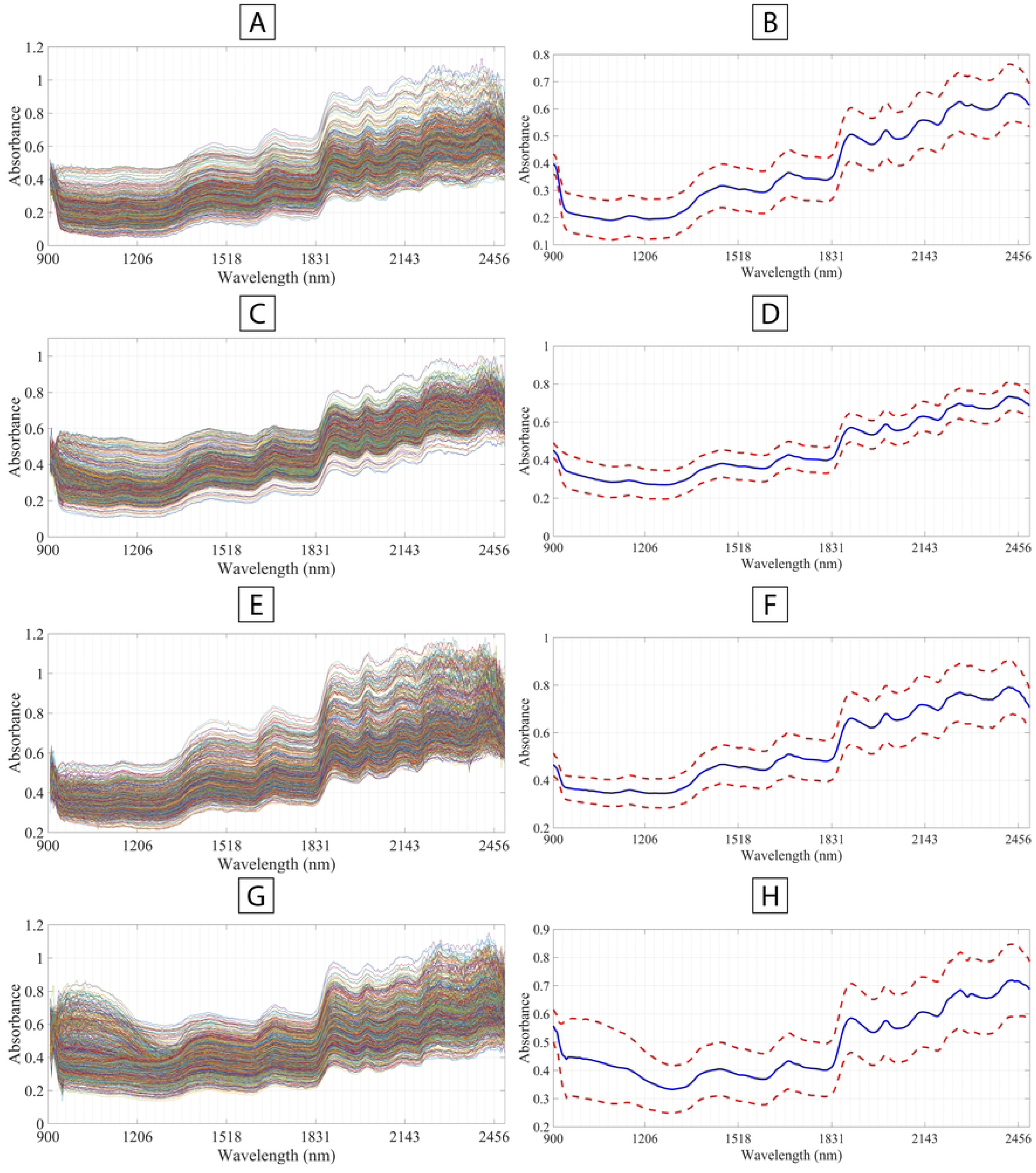
Unfolded (A, C, E, G, I) and mean spectra (B, D, F, H, J) obtained from the pronotum of four *Conocephalus* species. (A-B) *C. fasciatus*; (C-D) *C. versicolor*; (E-F) *C. iriodes*; (G-H) *C. saltator*. (B, D, F, H) red lines: standard deviation, blue line: mean spectra.

**Fig 5.**
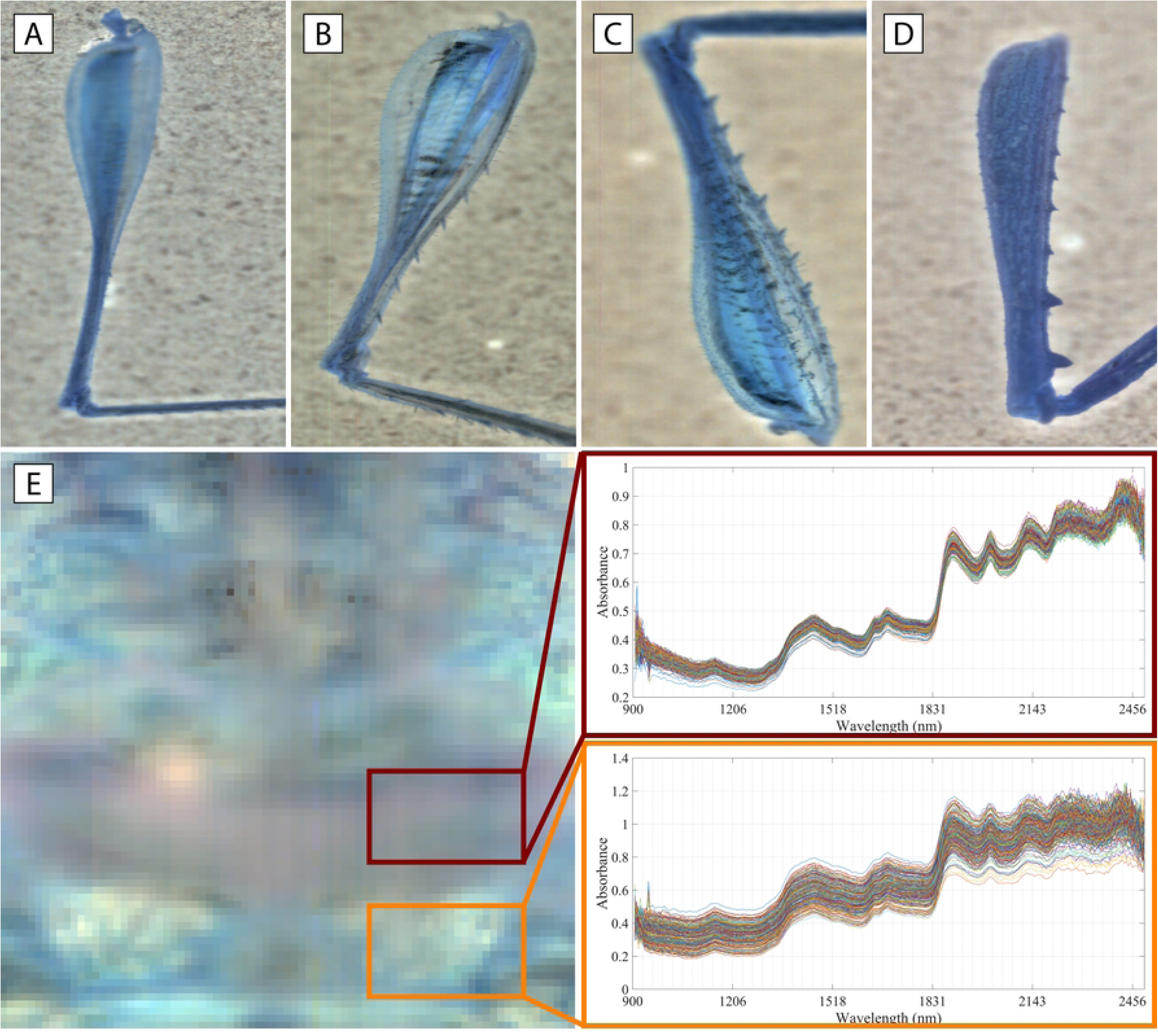
Transmitting capability of NIR radiation and spectra profile differentiation in katydid samples. False RGB image of legs of (A) a *Conocephalus fasciatus*, (B) a *Vestria viridis*, (C) a *Subria amazonica*, and (D) *a Typophyllum* sp.; (E) False RGB image of an *Acanthodis aquilina* pronotum with unfolded pre-processed spectra (threshold = 3, percentage zeros = 25) of two selected areas.

Another conspicuous band in our samples that is congruent with some previous authors [53,55,72,73] is between 1612 and 1831 nm (Figs 4C-E, 5A-J, 6G-H). This band may be attributed to functional groups found in the cuticular hydrocarbons, corresponding to the CH, CH_2_, or CH_3_ first overtones [40,46]. Other smaller bands attributed to hydrocarbons present in the cuticle and congruent with previous authors are between 1976 and 2064 nm (amide combination band of chitosan), 2064 and 2187 nm (C-C combination band), 2187 and 2360 nm (CH_2_ or CH_3_ combination band), and 2360 and 2500 nm (CH combination band) [46] (Tab 1, Figs 3C-E, 4A-H, and 5E). These bands vary in intensity among our samples and other insect taxa [53,55,72,73]; this characteristic and peak displacement information may be used as diagnostic characters.

An important spectral region in all our samples ranges between 900 and 1100 nm. Previous authors postulated that this region shows little valuable information, resulting from weak third overtones of fundamental absorbances [40,92] and, in our samples, it varied significantly in intensity among all the taxa studied (Figs 4A-H, 5E) —for example, in *Conocephalus* spp., absorptions in this range show an important variability (Fig 4A-H). Although this region is very noisy, it can be used as a diagnostic range for species after applying denoise pre-processing methods, such as SNV, derivate polynomials, or spatial binning.

Even though the literature postulates that NIRS equipment profiles cuticular hydrocarbons [23,40,46], our findings indicate that NIR radiation can penetrate (transmittance) significantly into the exoskeleton, reaching the musculature and beyond. In nearly all of our false RGB images of the legs, the musculature was clearly visible (Figs. 3A and 5A-C). However, in specimens with thicker cuticles, the musculature was less noticeable (Fig. 5D). In smaller samples, such as the legs of *Conocephalus* spp. and *P. teres*, the radiation was capable of penetrating the inner surface of the femur, allowing it to reach the background (Fig. 5B). This characteristic suggests that the organic molecules responsible for the recovered absorbance bands were not present solely on the exoskeleton but also internally, and the penetration capability may depend on (but not exclusively) the cuticle thickness. However, it is hard to ensure how deep the radiation can reach in regions with a thicker exoskeleton, such as the head and pronotum. The relief of the cuticle in these body parts is not always visible either in false RGB or decomposed near-infrared hyperspectral images. In species with large body sizes, such as *Acanthodis aquilina* (Linnaeus, 1758), the exoskeleton surface is more easily differentiated (Fig 5E), and the spectra are generally less noisy than in small species (Fig 5G). However, in small species, such as *Conocephalus* spp. or *Phlugis teres* (De Geer, 1773), the images revealed structures that do not correspond to the pronotal surface in many regions of the NIR range (S2 Fig), suggesting that the thicker cuticle of large specimens’ pronotum somehow limits the radiation from penetrating deeper.

The highest transmittance occurred in the wavelength region from 2030 to 2493.75 nm, corresponding to the longer wavelengths within the NIR spectral range. As mentioned before, this is one of the noisier regions in our images. We believe that variations in composition and physical characteristics across the surface, as well as within different parts of the sample (e.g., exoskeleton, muscles, and the surface of the femur), are likely contributing to the scattering of NIR light and the introduction of noise into the spectra in these regions.

The capacity to penetrate the insect body suggests that the spectra acquired by many previous authors [39,40,86,43–49,51] also correspond to the internal body’s composition, such as musculature, fluids, or organs. As evidenced here, how deep the radiation can penetrate may depend on the thickness and hardness of the sample and, therefore, may be influenced by the size of the specimens. In other words, the thinner and softer an insect’s cuticle, the easier it will be penetrated by NIR radiation.

Establishing the focus may be challenging when imaging insects. Due to the transmittance capacity of near-infrared radiation, the focus also may influence which information the imaging is accessing. As seen in Fig 3, the spectra of the same selected pixels in slightly different focus depths recover different spectra profiles (i.e., different chemical information). In area 1, where there are the muscles, the spectra profile remains more stable, suggesting that the chemical information recovered is also from the musculature, and the NIR radiation could not pass through it. On the other hand, the spectra of areas 2 and 3 varied more in intensity and noise, suggesting that different information is being accessed.

In Rodríguez-Fernández et al. [23], the authors used NIRS to distinguish nine species of flies of the genus *Neodexiopsis* Malloch. These are very small flies, 2-5 mm long. Due to the tiny size and the softer cuticle of these flies, we believe that the spectral profiles obtained by the authors do not correspond solely to cuticular hydrocarbon but also to the internal components. Grace et al. [51] also reported differences in the lipids expressed on the pronotal cuticle of eight populations of the grasshopper *Hesperotettix viridis* (Thomas), fed on two different host plants. As in the previous case, the results may not correspond solely to metabolic lipids expressed on the cuticle but also to organic molecules dispersed on the body’s internal components, such as organs or hemolymph. In Sovano et al. [44], the authors obtained spectra from the right anterior leg of dehydrated katydid specimens belonging to the genus *Aganacris*. This leg is much smaller than the hind legs, with a much thinner cuticle. We can assume that the radiation could pass through the forelegs much easier than the hind ones. However, the authors did not provide the spectral profiles, so it was not possible to corroborate or refute this hypothesis.

As the thorax has a thicker exoskeleton and denser thoracic musculature, resulting in less noisy spectra than legs, we assume that the former prevents deeper penetration of near-infrared radiation. Nonetheless, if the leg’s images are treated and pre-processed, they can still be used. According to the literature, spatial binning is an excellent option to reduce spectral noise and the size of the image, but it affects spatial resolution [79,93]. Spatial resolution was not an issue in our study because of the sample size and macro lens utilized for image acquisition. However, spatial binning must be used with caution in very small samples once the number of representative pixels is low. For leg image acquisition, we suggest that the images be taken directly on the reference Teflon plate to avoid interference from the styrofoam used here.

Leg spectra (Figs 3C-E) have more discrete absorbance bands than the pronotum (Figs 4A-H). This characteristic may be a consequence of the cuticle’s different composition in distinct body parts and may also (at some point) be influenced by the NIR penetration capability. When comparing images of the legs and body dorsum, it is possible to see that the spectra profiles are distinct. As seen in Figs 6A-D, when a PCA is performed comparing the dorsum of the head, pronotum, and stridulatory apparatus versus the proximal portion of the hind legs of a *C. fasciatus* specimen (for instance), it is possible to see that both parts grouped separately (Figs 6A-B). When analyzing the scores of the PCA, representative pixels of the body are grouped on the positive side of PC1, while only a few pixels of the leg are grouped on the same side (Figs 6C-D). The loadings of PC1 suggest that at least four spectral regions are primarily responsible for this behavior: approximately 1800 and 1850 nm, 1950 and 2000 nm, 1650 nm, and 2100 and 2110 nm (Fig 6E). PC2 recovered only intraclass differences. These results support the hypothesis that different body parts have distinct spectra profiles and only homologous body parts should be used in a comparative multitaxa analysis.

**Fig 6.**
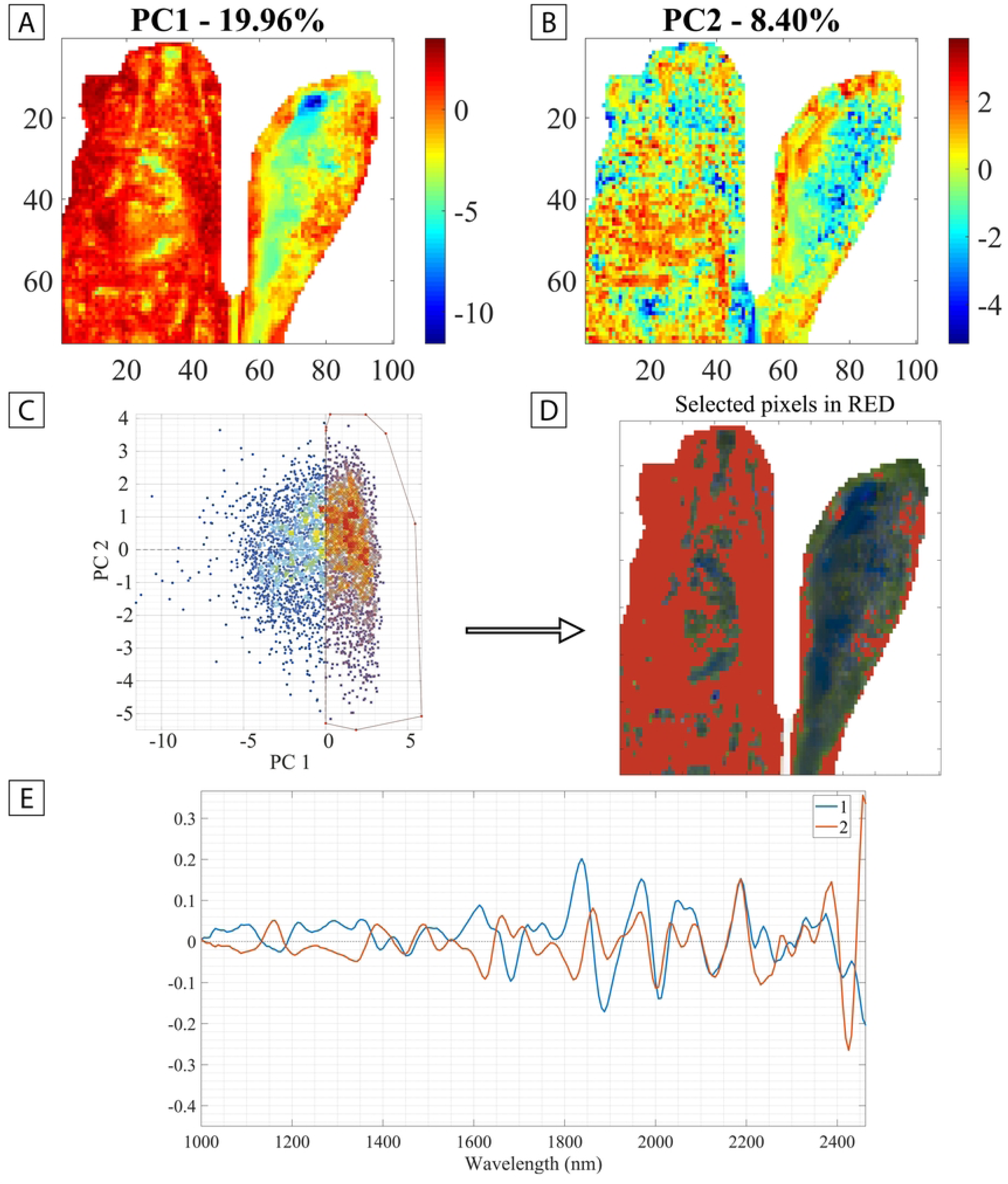
Comparison of the dorsum of the head, pronotum, and stridulatory apparatus versus the proximal half of the hind leg of *Conocephalus fasciatus* using Principal Components Analysis (PCA). (A) PC1. (B) PC2. Color temperature: signal intensity. (C) Scores for PC1 and PC2 with selected pixels. (D) Selected pixels of Fig C. (E) Loadings for PC1 and PC2.

Our findings demonstrate the highly heterogeneous nature of cuticles, wherein individual body appendages, structures, or regions may exhibit varied spectral profiles within the same sample (Figs 5E). The body and hind legs also have different spectra profiles, with some areas displaying very distinct spectra, which means they are also dissimilar regarding chemical composition (Fig 6). If we consider only the cuticle, these results corroborate with Moussian [52], who postulated that different body parts have different cuticle chemical and structural compositions.

### Sample pre-processing, dead pixels, and spikes

Our raw images present a considerable abundance of spikes, constituting one of the primary challenges we addressed (Figs 2C and D). We observed lines of pixels with spikes, and certain spectral regions consistently exhibited spikes (e.g., 943.75 nm, 975 nm, 1287.5 nm, 1912.5 nm, 2187.5 nm, 2200 nm, 2250 nm). The first case was probably caused by burnt or defective pixels in the camera sensor, and malfunctioning channels caused the second. These instrumental problems can occur and are common, and the solutions to deal with them are available in Hypertools and other software for hyperspectral image analysis [53,79].

Manual inspection and elimination of spikes can be performed, but it is too time-consuming. One of the most popular ways of automatically detecting them is using neighboring filters. Once detected, the value of the spike is automatically replaced by the median of the adjacent spectral channels [79]. Threshold selection needs optimization, and the signal-to-noise ratio must be considered [79]. Moboraki and Amigo [53] recommended a threshold value of 6. However, in practice, finding the proper threshold will depend on the image analyzed once it, in turn, depends on the mean spectrum of the image (and not necessarily the sample pictured) [79,94,95]. In other words, different thresholds may be applied to the same specimen depending on the images resulting from the scanning process. The noisier and more heterogeneous the spectra of a given image, the larger the standard deviation for the absorbance values over the mean spectrum; consequently, smaller thresholds will be necessary. Highly heterogeneous samples or images in which the background is an important part of its composition (e.g., images of legs) can also negatively affect the efficiency of the neighboring filters because these characteristics enlarge the standard deviation of the mean spectrum [79].

As some of the samples’ spectra are very noisy (especially those of the legs), the threshold applied for effectively identifying and eliminating spikes had to be adapted to each image. In some cropped images of the pronotum, a factor of four times the standard deviation of the mean spectrum was enough to eliminate the spikes. Still, in others, a lower value was necessary. For example, Figs 7A-F shows the thresholds tested for an image of a *C. fasciatus* pronotum. In this case, the threshold that best detected the spikes was a factor of two (Figs 7E and 8E). When considering a low threshold factor, some pixels containing scatter spectra are also identified as spikes, and a median spectrum substituted the entire spectrum of these pixels (Figs 7E and 8E). On the other hand, threshold factors from three to six could not satisfactorily identify and eliminate the spikes in this sample (Figs 7A-D and 8A-D).

**Fig 7.**
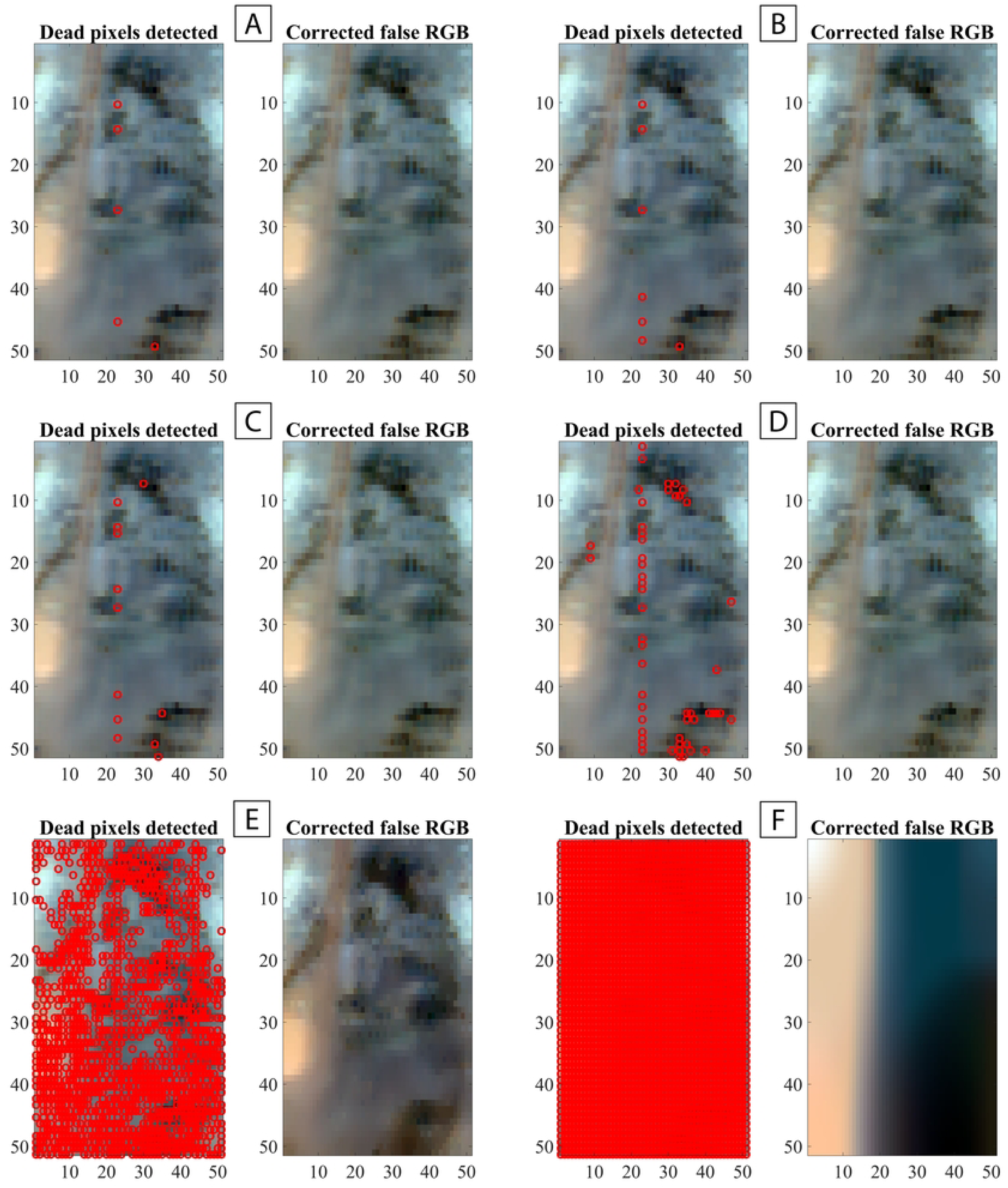
Evaluation of the efficiency of the neighboring filter to detect and eliminate spikes of a *Conocephalus fasciatus* pronotum image using different thresholds. (A) Threshold = 6. (B) Threshold = 5. (C) Threshold = 4. (D) Threshold = 3. (E) Threshold =2. (F) Threshold = 1.

**Fig 8.**
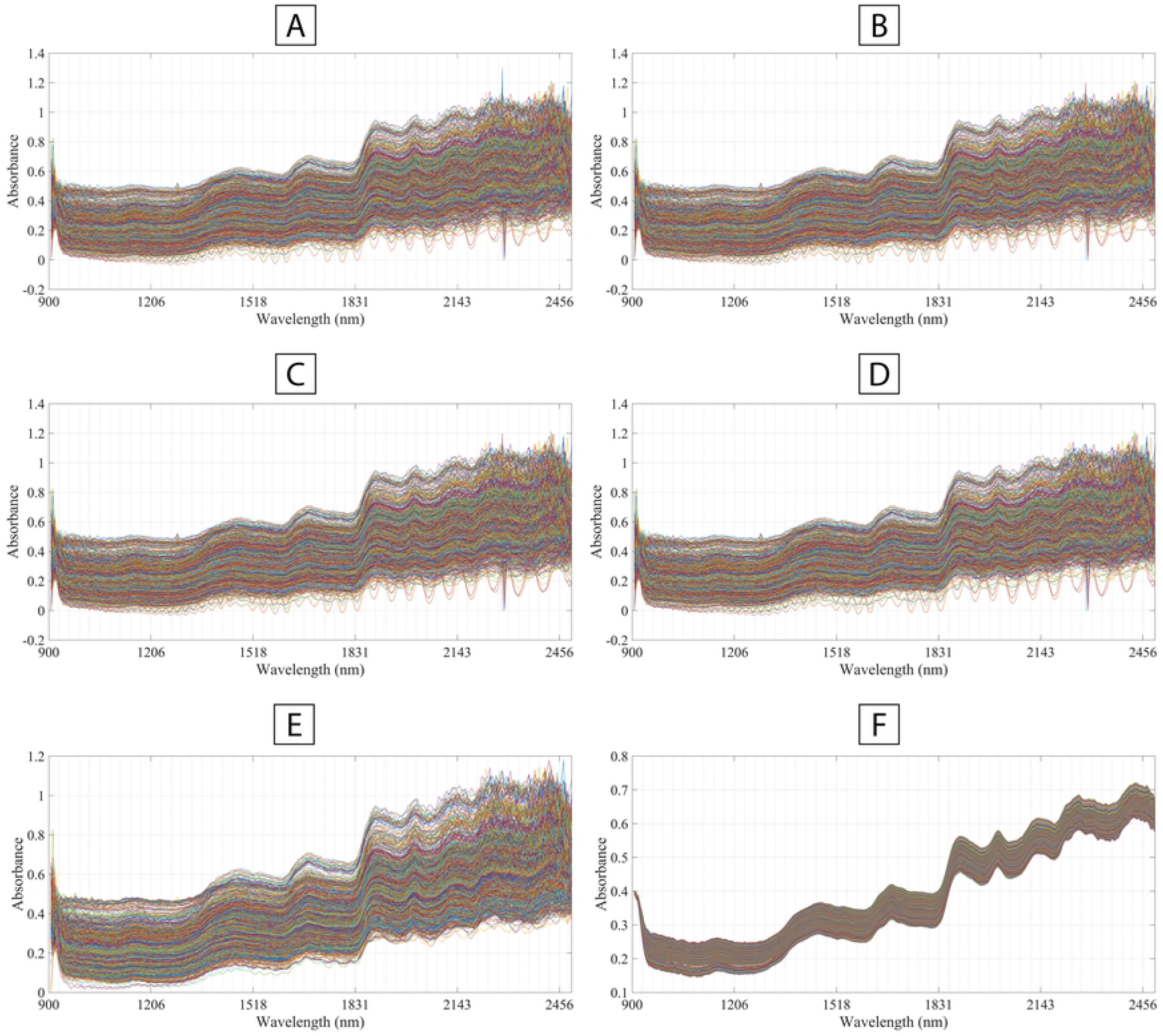
Unfolded spectra of a *Conocephalus fasciatus* pronotum image after applying the neighboring filter to detect and eliminate spikes using different thresholds. (A) Threshold = 6. (B) Threshold = 5. (C) Threshold = 4. (D) Threshold = 3. (E) Threshold =2. (F) Threshold = 1.

In some cases, the standard deviation of the mean spectrum was too large, and even a threshold value of two was not enough to detect some very large spikes. These spikes negatively affected our analysis, specifically the PCA, once they were not removed even after the Savitzky-Golay polynomial smoothing. In all tested images, a threshold factor of one detected all the spectra and completely replaced them with median spectra, thereby modifying all the information in the hypercubes (Figs 7F; 8F). Hence, it should be avoided. When even a threshold value of two remains unsuitable, performing spatial binning can be a viable solution to remove spikes, resulting in substantial spectral denoising. Manual detection and elimination can be made if some spike still remains. Neighboring filters may be time-consuming, depending on the size of the image (i.e., the number of pixels), the signal-to-noise ratio, optimization of the threshold factor, and the computer’s processing capability [79].

Savitzky-Golay polynomial derivative and smoothing filter with a first-order polynomial (2^nd^ degree) were evaluated using window sizes varying from 9 to 21 points, aiming to enhance the signal-to-noise ratio and minimize light scattering effects. Larger window sizes (from 17 to 21 points) were found to be inefficient due to the limited spectral resolution (only 256 spectral channels). Furthermore, a pronounced border effect was noted, resulting in a "tail" of points at the spectral ends. [96]. As previously discussed, relevant information appears to be present at the ends of our sample spectra, especially for interclass differentiation. Consequently, window sizes exceeding 16 were avoided. Conversely, window sizes varying from 9 to 15 demonstrated better suitability. In exploratory analysis, these window sizes exhibited minimal variance in intraclass differentiation. However, the window size of 15 points yielded the highest percentage of maximum variance for the first two principal components (Fig 10A-B) and was therefore chosen as the optimal window size for the second set of images. The Savitzky-Golay polynomial derivative filter was applied prior to Standard Normal Variate (SNV) processing as it enhanced interclass differentiation (Fig. 9).

**Fig. 9.**
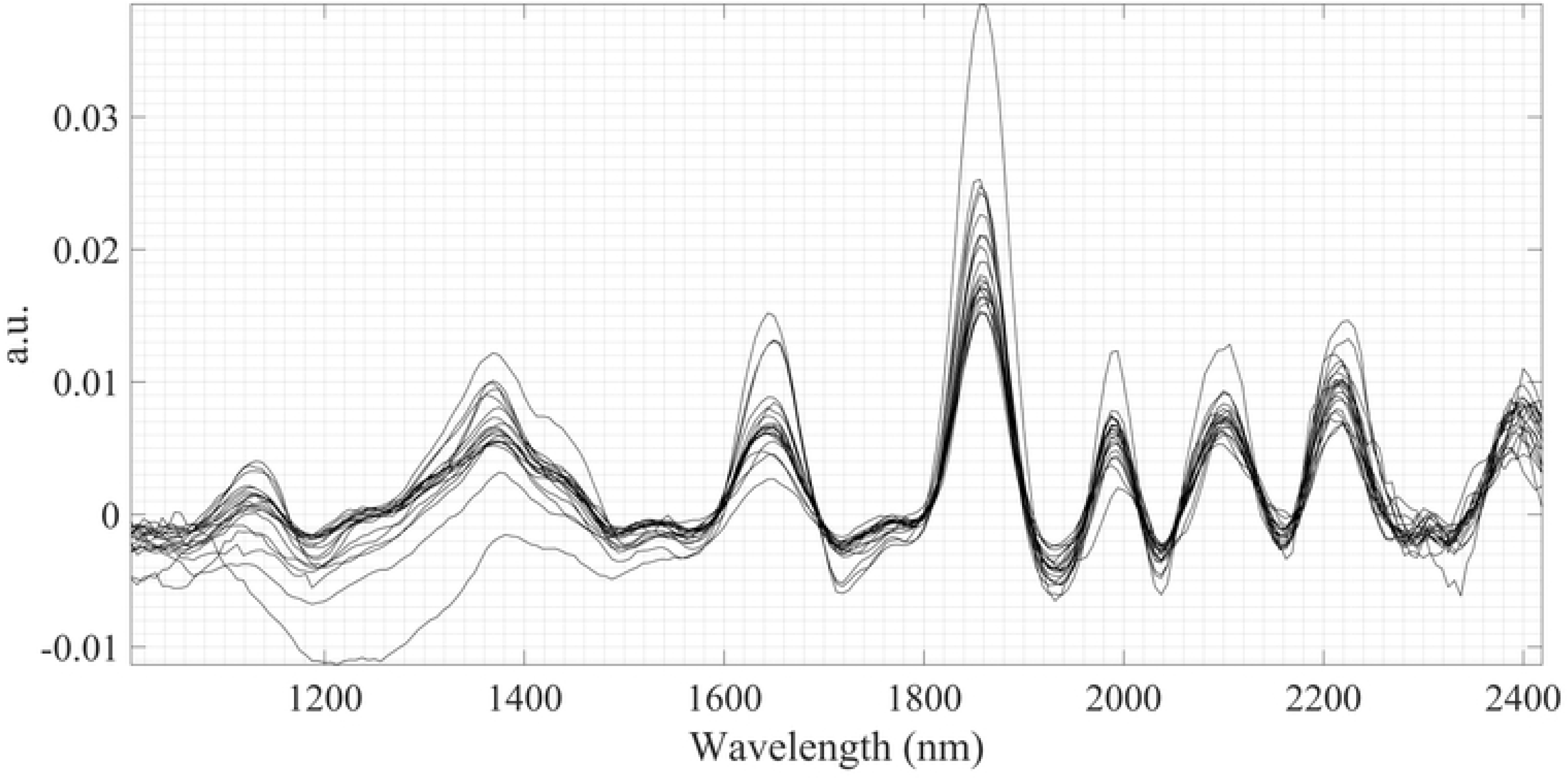
*Conocephalus* species’ 20-most representative pre-processed spectra with Savitzky-Golay polynomial derivative and smoothing filter with a first-order polynomial (2^nd^ degree and 15 points window) and Standard Normal Variate (SNV). a.u. = arbitrary unit.

**Fig 10.**
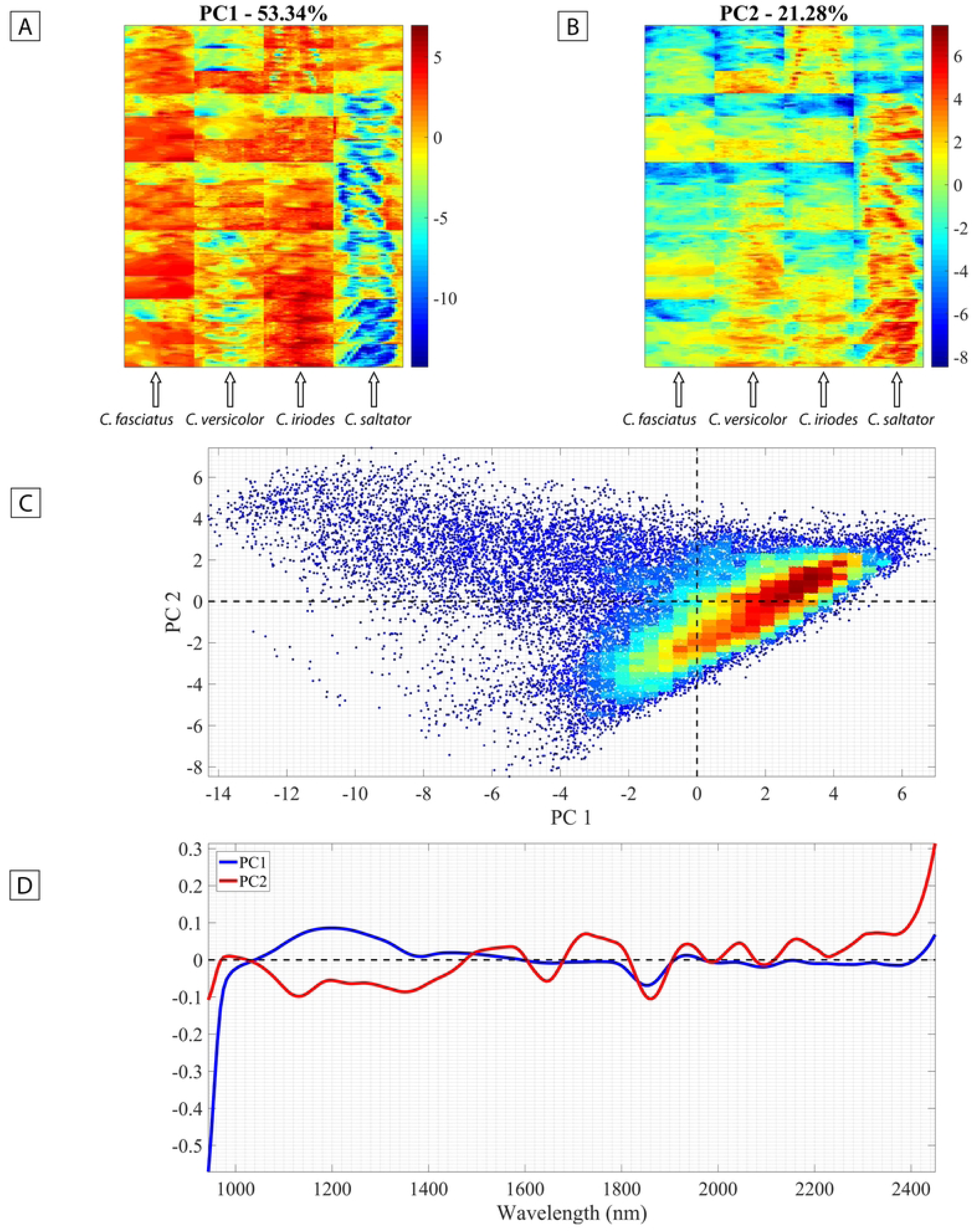
Principal Components Analysis (PCA) results. (A) PC1; (B) PC2 (color temperature = signal intensity); (C) Scores plot of PC1 x PC2 (color temperature = pixels concentration); (D) Loadings for PC1 and PC2.

With the window size of 15 points for the Savitzky-Golay polynomial derivative chosen for the second set of samples, the spectra ends were cropped to result in the final NIR range from 944 to 2450 nm. Despite previous authors postulating that wavelengths below 1000 nm show limited information due to weak third overtones absorptions [40,92], we decided to use the maximum range possible based on significant visual differences observed in this region within the images studied. The threshold values most suitable for spike removal, in this case, were 2 and 3.

Ultimately, spatial binning is effective for improving the quality of spectral information. However, when applied to small samples or low-resolution images, researchers should carefully consider the window size (i.e., the number of pixels to be combined). Larger windows can reduce spatial resolution, potentially masking spectral heterogeneity and intrinsic characteristics. Manual spike removal should be employed if necessary.

### Discriminating *Conocephalus* species

#### Principal Components Analysis (PCA)

A false RGB image and unfolded spectra profile for the triplicate sample of each species of *Conocephalus* used are available in S3 Fig. The PCs that recovered the pattern of interclass differentiation of the samples (species) were 1 and 2, each explaining 53.34% and 21.28% of the total data variation, respectively (Figs 10A and B). As seen in Figs 10C and D, the positive scores of PC1 are influenced by pixels with a more intense absorbance band between 1040 and 1360 nm; however, the negative scores are more strongly influenced by lower absorbance values at the beginning of the spectra, before 1020 nm (Fig 10D). This result condenses a representative part of the pixels of *C. fasciatus* and *C. iriodes* on the positive side of the distribution on the PC1 axis (Fig 10C).

On the other hand, the PC2 loadings reveal that pixels with intense absorbance values at the end of the spectra, after 2400 nm, strongly influence the distribution of scores for the positive side (Fig. 10E). Absorbance bands between 1480 and 1600, 1660 and 1820, 1900 and 1980, 2000 and 2080, 2120 and 2220 nm also influence the ordination of pixels on the positive side of PC2. Representative pixels of *C. versicolor*, *C. iriodes*, and *C. saltator* with intenser values in the aforementioned NIR regions tend to be condensed on the positive side of the PC2 axis (Fig 10B and C).

#### Partial Least Squares Discriminant Analysis (PLS-DA)

As mentioned before, five specimens of each species of *Conocephalus* (except *C. equatorialis*) were used to generate the PLS-DA classification model (S1 Table); the remaining specimens were used in the external validation (*C. fasciatus* n=5; *C. iriodes* n=3; *C. versicolor* n=4; *C. saltator* n=10). Twenty latent variables (LVs) were selected for the PLS-DA classification model based on the highest sensitivity and specificity values. In the cross-validation phase, 77% of the pixels were assigned to one of the four classes, and 23% remained unassigned (see Table 2 and Fig 11). An overall classification accuracy of 90% for the assigned pixels (error rate: 10%) was achieved, which indicated the effectiveness of the PLS-DA model in assigning the pixels to the correct classes (i.e., species) (Table 2-3 and Fig 11). As shown in Table 3 and Fig 11, class precision was high, with the lowest value at 87% for *C. iriodes* and the highest reaching 94% for *C. saltator.* Detailed results for each class can be seen in S4 and S5 Figs.

**Fig 11.**
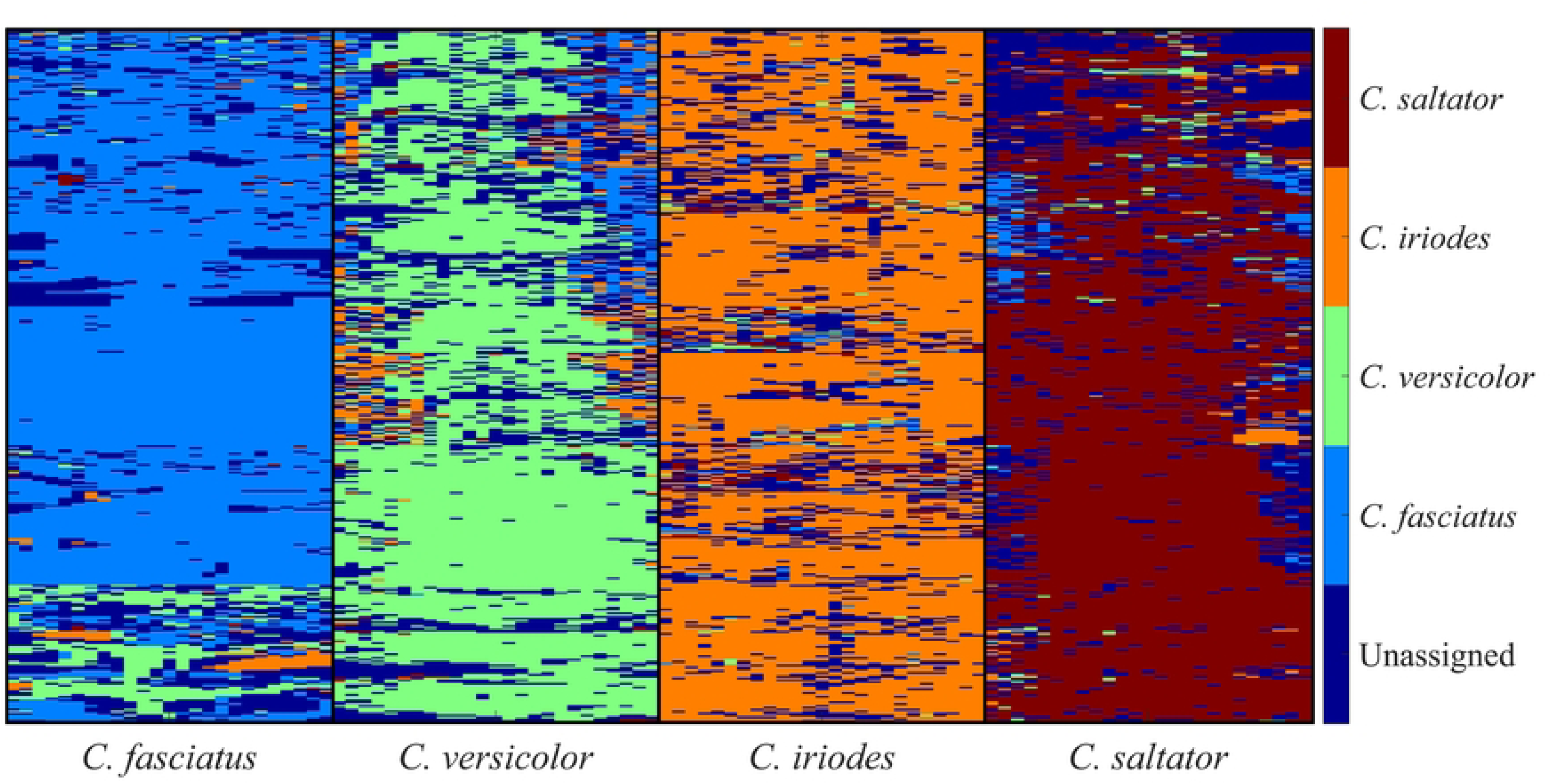
Partial Least-Squared Discriminant Analysis (PLS-DA) calibration and cross-validation results.

**Table 2.** Confusion matrix of the cross-validation model of PLS-DA. Values = number of pixels.

| Real/predicted | *C. fasciatus* | *C. versicolor* | *C. iriodes* | *C. saltator* | Unassigned |
| --- | --- | --- | --- | --- | --- |
| *C. fasciatus* | 6708 | 593 | 139 | 25 | 1910 |
| *C. versicolor* | 576 | 5551 | 500 | 113 | 2601 |
| *C. iriodes* | 121 | 96 | 6882 | 241 | 2013 |
| *C. saltator* | 217 | 136 | 196 | 6511 | 2261 |

**Table 3.** Merit figures for each class of the cross-validation model of PLS-DA.

| **Classes** | **Specificity** | **Sensitivity** | **Accuracy** |
| --- | --- | --- | --- |
| *C. fasciatus* | 96% | 90% | 88% |
| *C. versicolor* | 96% | 82% | 87% |
| *C. iriodes* | 96% | 94% | 89% |
| *C. saltator* | 98% | 92% | 94% |

The scores of the built PLS-DA model are presented in Fig 12A (LV1 x LV2) and Fig 12B (LV1 x LV3). The loadings of LV1 revealed that the spectral range below 1100 nm is critical for classifying *Conocephalus* species. Notably, the first three LVs were significantly influenced by the absorbance values between 900 and 1100 nm (Figs 13A-C). These results contrast with previous studies that mention that this region provides little valuable information due to weak third overtones absorptions [40,92]. Additionally, the loadings of LV2 and LV3 indicated a strong influence from absorptions beyond 2200 nm.

**Fig 12.**
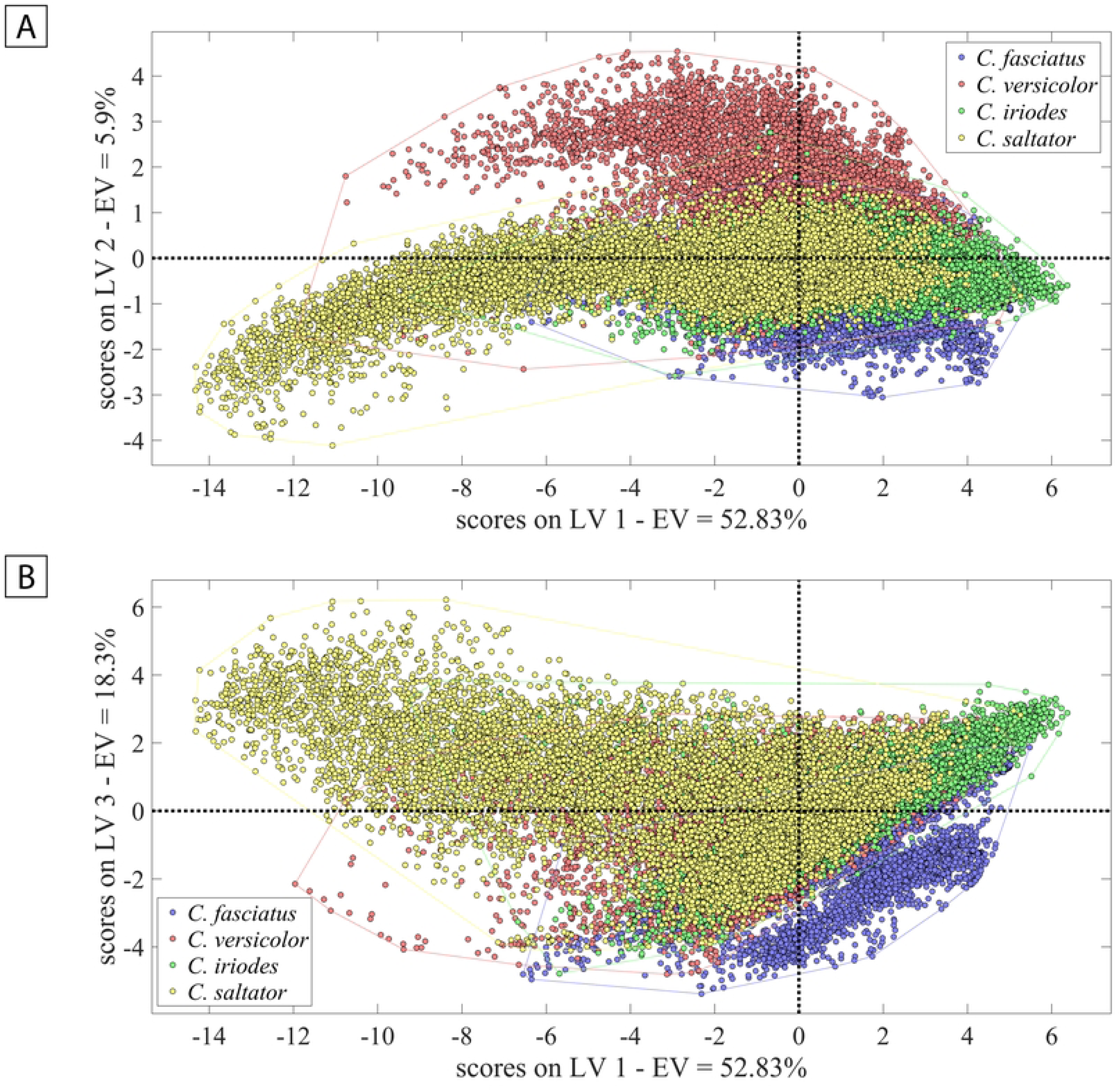
Scatter plot of the Partial Least-Squared Discriminant Analysis (PLS-DA) model. (A) First and second latent variables space. (B) First and second latent variables space.

**Fig 13.**
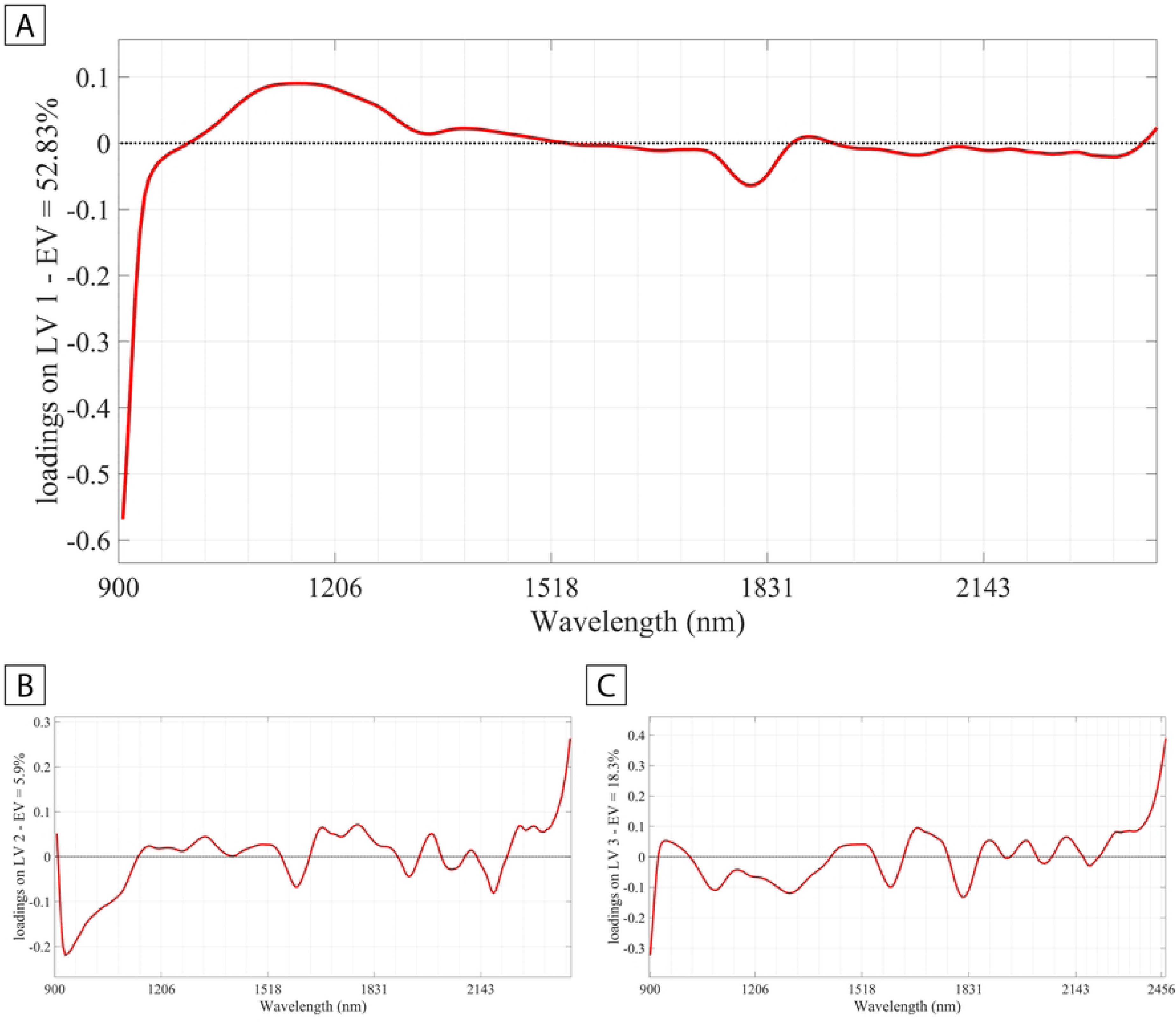
Partial Least-Squared Discriminant Analysis (PLS-DA) loadings results. (A) LV1. (B) LV2. (C) LV3.

The classification result for *C.* versicolor was the least accurate (sensibility=82%, accuracy=87%), with many pixels unassigned (n=2601, see Table 2 and S3 Table). As seen in the S3 Table, two specimens had all three replicates with partial classification success; one had two replicates with partial classification, and another replicate failed; one had one replicate with partial classification and two successes; and two had all three replicates successfully classified. It appears that some *C. versicolor* specimens were exposed to chemicals that altered their original colors, like ethyl acetate or alcohol, commonly used in sacrifice recipients and traps by entomologists. The exposition of these chemicals may have affected pixel classification. However, other specimens, apparently treated under the same condition and used to model the PLS-DA, achieved more accurate classification (S3 Table). Therefore, it remains unclear whether the misclassification was due to the usage of chemicals or imaging acquisition issues.

An external validation phase was conducted, and the PLS-DA model was applied to classify *Conocephalus* specimens that were not included in the modeling development phase. The model correctly assigned most pixels of each specimen to the correct class/species (Figs 14A-B, 15A-B, and S3 Table), even of *C. versicolor*, in which only one specimen had its triplicates partially classified (i.e., pixel counting ≥ 30% and < 60%; specimen 041). One specimen of *C. saltator* had one replicate misclassified (pixel counting < 30%) and two partially correct. When accounting for the number of specimens correctly classified, the model misclassified two specimens: one *C. fasciatus* (voucher 007) (Fig 14A, *dashed line*) and one *C. iriodes* (voucher 037) (Fig 14B, *dashed line*).

**Fig 14.**
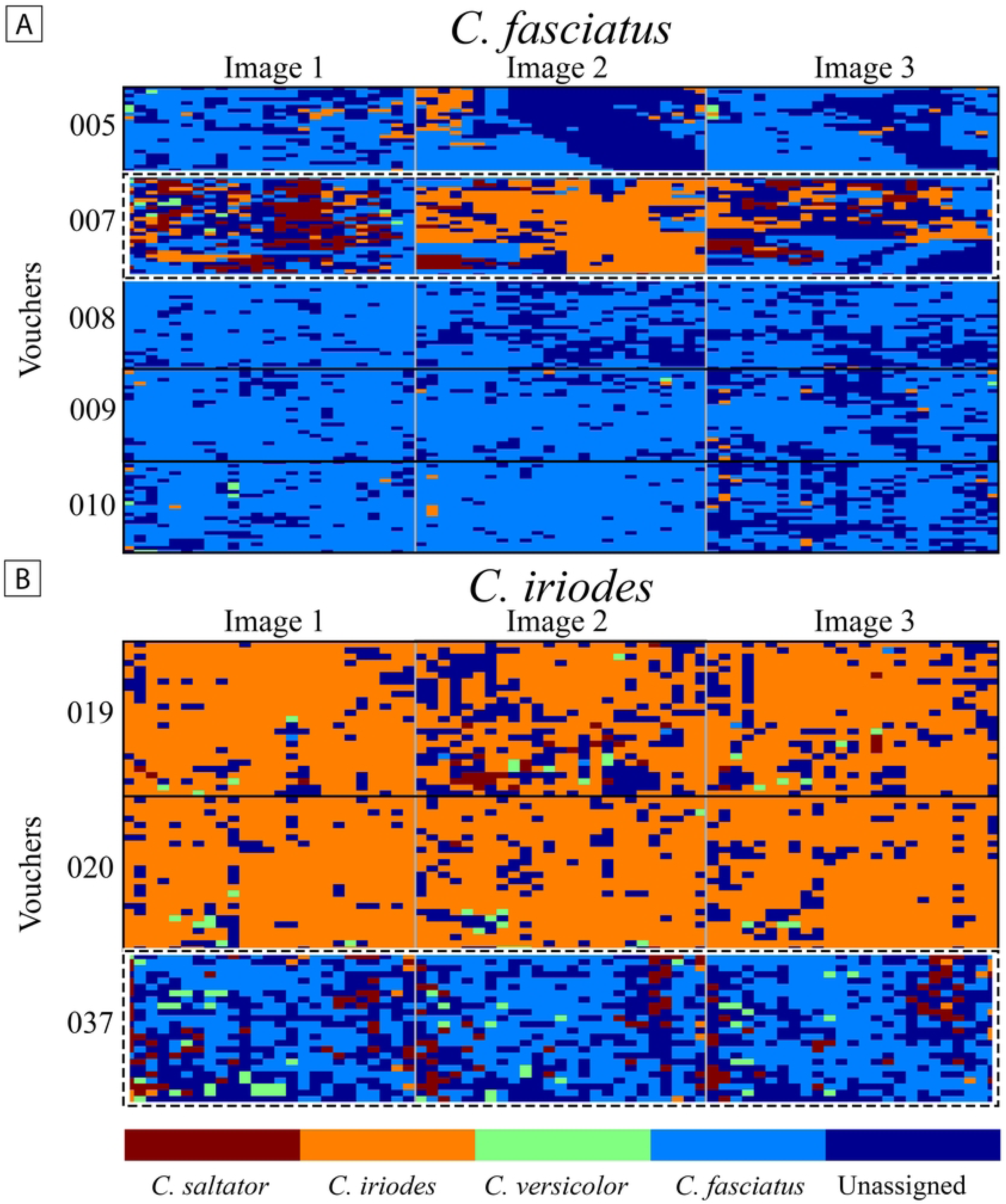
Prediction test of Partial Least-Squared Discriminant Analysis (PLS-DA) model on external validation set. (A) *Conocephalus fasciatus*. (B) *Conocephalus iriodes*. Dashed line = samples that failed.

Regarding the specimen of *C. fasciatus* (voucher 007), we have little doubt about its species identification. One replicate was still partially assigned to the correct species, but different aspects may have caused this misclassification of the other two replicates. We primarily attribute this to focus position issues. Imaging simultaneously multiple specimens led to an incorrect focus on the pronotum dorsal surface (see S6A Fig). This assumption is plausible given the differing pixel classification patterns among the replicates, with the third image showing better accuracy due to improved focus with a partial classification (Fig 14B, S6A Fig and S3 Table). The second image of voucher 007 performed the worst in pixel assignment among its replicates, which reinforces this hypothesis once it was the image most out of focus. The same case may be attributed to the last image of *C. saltator*’ specimen 025. This sole image was out of focus, and the other two images, which were in a better focus position, were partially assigned to the correct species (S3 Table).

The PLS-DA model also misclassified voucher 037 (Fig 14B), tentatively determined as *C.* cf. *iriodes.* Unlike the previous case, ensuring this female’s species identity is challenging. In fact, it may belong to a different species, as the pixel classification patterns were consistent across the three images, with most pixels incorrectly assigned to *C. fasciatus*. Additionally, focus adjustment was not an issue in this case, as all triplicated images were focused on the pronotum and head (S6B Fig). This is an optimistic result once it shows that the model was capable of indicating a misclassification.

*Conocephalus versicolor* and *C. saltator* are two polymorphic species bearing variations on the wing development and postabdomen appendages [75–78]. Specimens of the two species, of both sexes, collected at different sites, with variations on the structures mentioned above, were tested. Although the model for *C. versicolor* was the least accurate, the prediction results were satisfactory as most pixels of long- and short-winged specimens were correctly classified (Fig 15A). The model for *C. saltator* performed the best in the cross-validation stage, accurately classifying most pixels for the specimens. In the external validation phase, it also performed very well despite the problem with the focus position in one replicate. We may assume that, for this species, the model was satisfactory, regardless of the polymorphism (Fig 15B).

**Fig 15.**
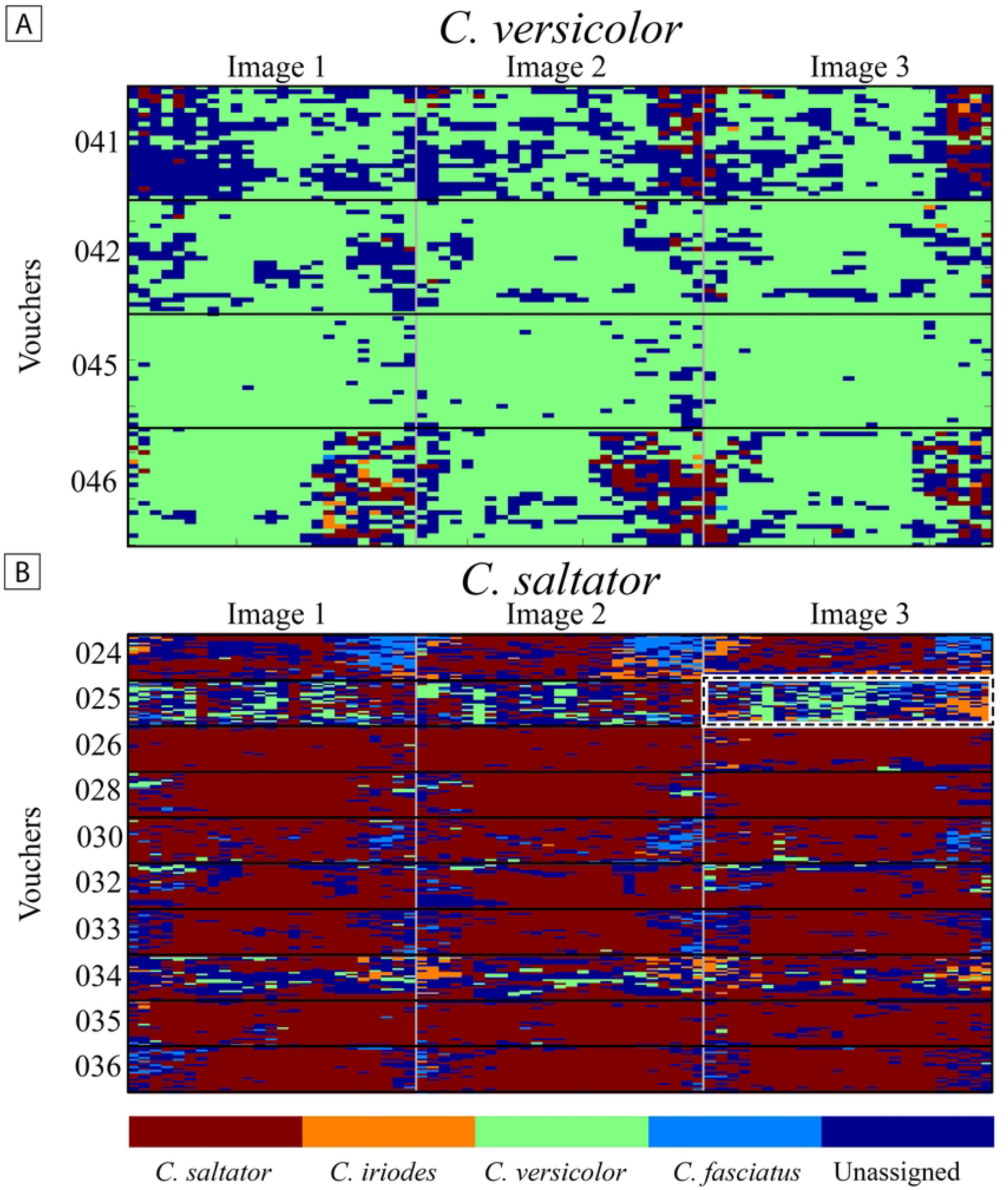
Prediction test of Partial Least-Squared Discriminant Analysis (PLS-DA) model on external validation set. (A) *Conocephalus versicolor*. (B) *Conocephalus saltator*.

Finally, the built PLS-DA model was applied to four specimens of *Conocephalus equatorialis*. As a result, the pixel classification pattern was inconsistent, and no evident classification pattern was noticeable (Fig 16 and S3 Table). This result is consistent with the fact that no specimen of this species was included in the development phase of the PLS-DA model, and the model failed to classify this species.

**Fig 16.**
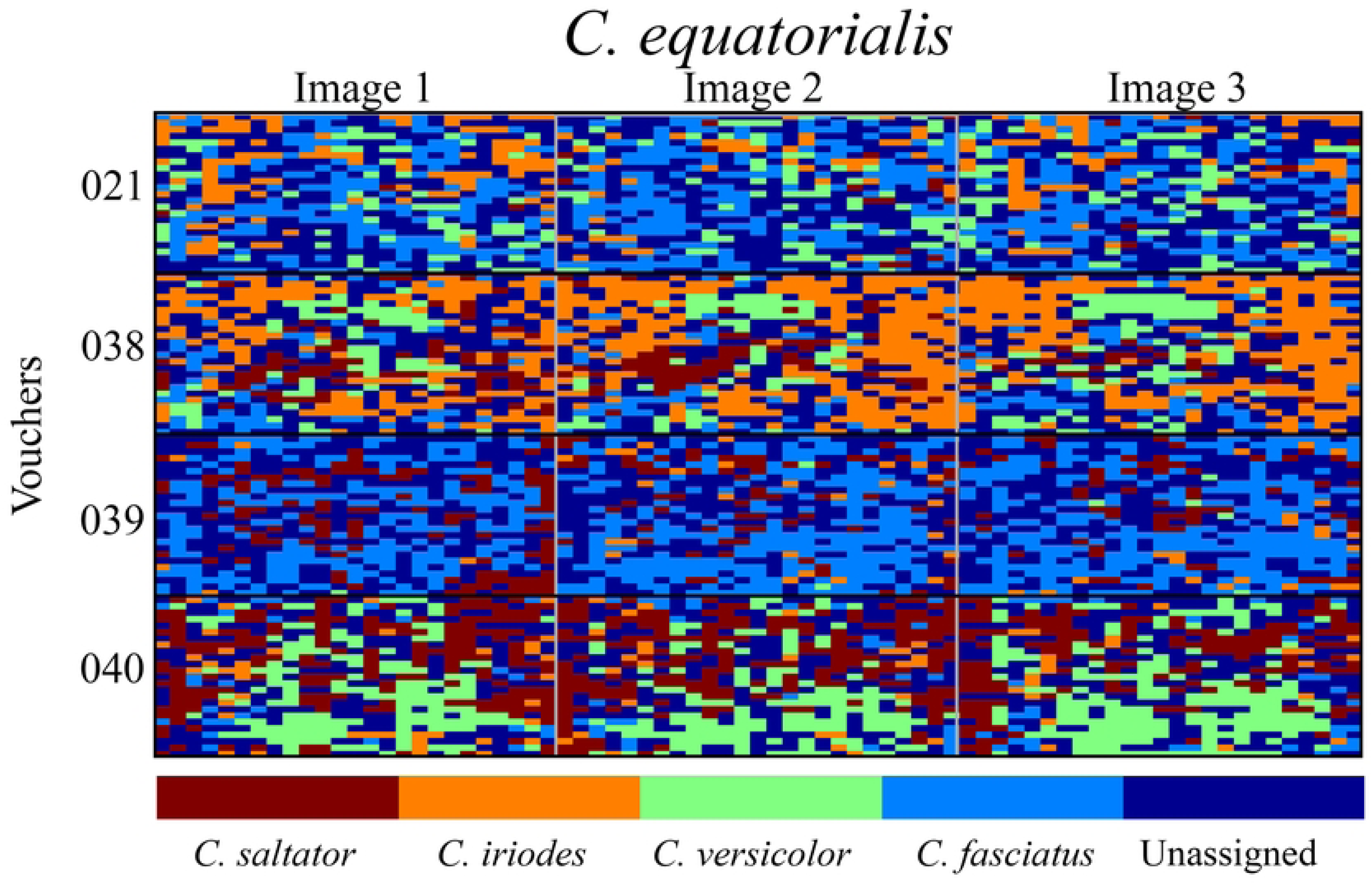
Prediction text of Partial Least-Squared Discriminant Analysis (PLS-DA) model tested in *Conocephalus equatorialis* specimens.

## Final considerations

HIS-NIR is a true nondestructive method that preserves the integrity of the samples. It is a suitable method for dried museum specimens, including types, once we have tested on old, new, clean, and dirty specimens. Furthermore, the acquisition method proposed here may be one of the best choices for taxonomic purposes [i.e., characterizing and differentiating taxa]. Setting up an image acquisition protocol facilitates future authors’ comparison of data.

Our study showed that images from different body parts [pronotum and legs] generated different spectra due to the nature of the exoskeleton and internal components. Therefore, we encourage that only homologous parts be used to model and compare species to decrease the likelihood of misclassification resulting from non-homologous information. Additionally, issues like transmittance can affect results. For example, leg images in this study were problematic, as radiation sometimes penetrated through the legs, capturing information from the background material. Standardizing the background material, such as Teflon, might help to mitigate this issue. Given the importance of comparing homologous regions and the heterogeneity of samples, this technology can also be explored in evolutionary studies by providing a new line of characters to be included in a phylogenetic matrix.

On the other hand, establishing a single spectra pre-processing protocol can be challenging and perhaps not beneficial to a taxonomic perspective since the pre-processing used will vary with the signal quality obtained. The signal quality (i.e., the presence of noise, scattering, dead pixels, and spikes) may vary according to the equipment used and the sample. Equipment produced by different companies operates in distinct spectral and spatial resolutions and different spectral bands and ranges. Differences in spectral dimension between equipment are the most concerning issue to the comprehensive appliance of the method because a calibration transfer or standardization of equipment for calibration transfer used by taxonomists would be necessary. Standardization in calibration transfer methods and equipment choice for calibration are subjects that must be discussed more in biodiversity sciences using NIR spectroscopy. However, standard equipment choice must also be cautiously addressed once it can promote scientific inequality and domain by groups with more financial support (e.g., from developed countries) concentrating the decision-making.

Another critical aspect to consider for the widespread adoption of hyperspectral imaging and global near-infrared spectroscopy in taxonomy is ensuring data accessibility for other researchers. Making this data available allows for the verification or refutation of taxonomic decisions supported by these methods and facilitates accurate identification by other scientists without needing to generate new classification models. However, a significant challenge with hyperspectral images is their file size. A practical solution is to archive RAW images and their metadata in permanent online repositories similar to those used in genomics (e.g., GenBank, BOLD, and NCBI). Establishing standardized protocols that align with good taxonomic practices will also be essential for effective data sharing.

Searching for chemical particularities encoded in spectral signatures all over the body and appendages of a given taxon is a new perspective for the integrative taxonomy of insects. The selected areas can be analyzed at least from two perspectives: (1) by studying each pixel’s spectrum, searching for unique signature patterns present in the sample and considering its heterogeneity, or (2) by the average spectrum of all pixels. A spectra library can be created with the results, and classification algorithms can be calibrated to determine future samples according to their spectra profile.

The usage of metabolomics and spectroscopical techniques in animal taxonomy is still an incipient field. However, our work is the first to use almost the entire near-infrared range and the first to use hyperspectral imaging technology for Orthoptera taxonomy. The discriminant model generated by the PLS-DA, with an assigned pixels’ accuracy of 90% for the species of the *Conocephalus*, showed the potential of hyperspectral imaging technology coupled with near-infrared spectroscopy to discriminate species. It is crucial to remember that no variable selection method (used to eliminate non-informative spectral information to the model) was performed prior to the modeling stage. Variable selection methods may be an excellent option for taxonomic purposes because they increase the model’s accuracy. Nonetheless, the results presented here are promising for insect taxonomy, as complementary and integrative methods are becoming increasingly essential. The approach demonstrated in this study effectively distinguished between species, underscoring its potential utility in this field.

## Acknowledgments

We extend our sincere gratitude to Embrapa Algodão, with special thanks to Dr. Everaldo Medeiros and laboratory technician Joabson Borges de Araújo for their generosity in providing equipment and invaluable support throughout this work. This study was funded in part by the Conselho Nacional de Desenvolvimento Científico e Tecnológico (CNPq), Fundação Coordenação de Aperfeiçoamento de Pessoal de Nível Superior (CAPES), the Fundação Amazônia de Amparo a Estudos e Pesquisas (FAPESPA), and the Programa de Apoio à Publicação Qualificada (PAPQ) of the Universidade Federal do Pará (UFPA)—GC Tavares: CNPq 421693/2022-3, CAPES 88882.444627/2019-01, and FAPESPA 2023/157870; JAM Fernandes: CNPq 310436/2021-4. GC Tavares also wishes to thank the Orthopterists’ Society for the financial support provided by the OSF Grant, which enabled the acquisition of equipment that greatly enhanced the quality of this research.

## Supporting information

**S1 Fig. Hyperspectral images acquisition.** (A-B) Orthoptera specimens during the scanning process with the Hyperspectral Imaging Spectrometer. (C-D) Specimens on the styrofoam plates.

**S2 Fig. Penetration capability of NIR spectra on a *Conocephalus fasciatus* specimen.** (A) Head, pronotum, and stridulatory apparatus. (B) Proximal half of hind leg.

**S3 Fig. False RGB images and spectra of *Conocephalus* species pronotum used to calibrate the Partial Least Squares-Discriminant Analysis (PLS-DA) model.**

**S4 Fig. Partial Least Squares-Discriminant Analysis (PLS-DA) cross-validation response for *Conocephalus* species.** (A) *C. fasciatus*. (B) *C. versicolor*.

**S5 Fig. Partial Least Squares-Discriminant Analysis (PLS-DA) cross-validation response for *Conocephalus* species.** (A) *C. iriodes*. (B) *C. saltator*.

**S6 Fig. Triplicates of false RGB images of *Conocephalus* specimens that the Partial Least Squares-Discriminant Analysis (PLS-DA) model failed in correctly classifying.** (A) *C. fasciatus*, voucher 021. (B) *C.* cf. *iriodes*, voucher 037.

**S1 Table. List of Orthoptera specimens used.** Underlined vouchers = specimens used in the PLS-DA modeling stage.

**S2 Table. List of images made of Orthoptera specimens with temperature and humidity measured at the moment of the scanning.**

**S3 Table. Percentage of pixel assignment per voucher image of Conocephalus.** Underlined data = specimens used in the PLS-DA modeling stage.

## Notes

### Competing Interest Statement

The authors have declared no competing interest.

